# Computationally designed stem-epitope mimetics elicit broadly reactive antibodies

**DOI:** 10.1101/2025.01.22.634229

**Authors:** Sarah Wehrle, Andreas Scheck, Laura Reusch, Flavio Matassoli, Sandrine Georgeon, Karla M. Castro, Johannes Cramer, Wayne Harshbarger, Stéphane Rosset, Sarah F. Andrews, Karin Schön, Badiaa Bouzya, Ronan Rouxel, Normand Blais, Enrico Malito, Adrian McDermott, Thomas Krey, Corey P. Mallett, Ventzislav Vassilev, Davide Angeletti, Bruno E. Correia

**Affiliations:** Institute of Bioengineering, École Polytechnique Fédérale de Lausanne, Lausanne CH-1015, Switzerland; Swiss Institute of Bioinformatics (SIB), Lausanne CH-1015, Switzerland; Department of Microbiology and Immunology, University of Gothenburg, Gothenburg, Sweden; Vaccine Research Center, National Institute of Allergy and Infectious Diseases, National Institutes of Health, Bethesda, MD, USA; GSK, Rockville, MD, USA; GSK, Rixensart, Belgium; Institute of Virology, Hannover Medical School, Hannover 30625, Germany; Center of Structural and Cell Biology in Medicine, Institute of Biochemistry, University of Lübeck, Lübeck, Germany; German Center for Infection Research (DZIF), partner site Hamburg-Lübeck-Borstel-Riems, Germany; Cluster of Excellence RESIST (EXC 2155), Hannover Medical School, Hannover, Germany; Centre for Structural Systems Biology (CSSB), Hamburg, Germany

**Author notes:** Correspondence (D.A.), (B.E.C.). These authors contributed equally (Sarah Wehrle, Andreas Scheck, Laura Reusch). These authors contributed equally (Davide Angeletti, Bruno E. Correia).

**Keywords:** computational protein design, epitope-focused vaccines, influenza virus, hemagglutinin, broadly neutralizing antibodies, surface mimicry

## Abstract

Broad protection against diverse influenza viruses can be conferred by broadly neutralizing antibodies (bnAbs) targeting a conserved site on the hemagglutinin (HA) stem domain. However, the low immunogenicity of this antigenic region hinders the robust induction of such antibodies. Here, we showcase a structure-based immunogen design strategy focusing on the surface mimicry of antigenic sites. By leveraging the structural definition of a stem epitope, we apply computational protein design to develop epitope mimetics to focus the immune response against this site of viral vulnerability. The structurally complex antigenic site is displayed on heterologous protein scaffolds, retaining excellent binding towards known HA stem-specific bnAbs. Our epitope-mimetic induces stem-specific antibodies against highly divergent group 1 and 2 subtypes. The results provide a general framework for the design of novel immunogens eliciting focused immune responses which may be a valuable tool in the development of effective vaccine candidates against other variable pathogens.

## Introduction

Influenza remains a major public health threat, causing severe morbidity in 3-5 million patients and estimated up to 650,000 deaths every year (Lee et al. 2010; Iuliano et al. 2018). Current vaccination strategies heavily rely on annual predictions of circulating strains to produce vaccines for the upcoming season and need yearly reformulation. However, vaccine effectiveness is often poor, ranging from 10% to 60%, due to vaccine mismatch (CDC 2020). This further highlights our limitations in pandemic preparedness, as novel strains emerging from zoonotic reservoirs remain a constant threat (Kanekiyo and Graham 2020; Yamayoshi and Kawaoka 2019; Neumann, Noda, and Kawaoka 2009).

Influenza vaccines are most commonly manufactured in eggs as either live-attenuated or inactivated-virus formulations. However, growth in eggs can lead to egg-adapted mutations that decrease immunogenicity (Chen, Zhou, and Jin 2010; Raymond et al. 2016). Influenza immunity is further complicated by immunodominance hierarchies as the main immune response is mounted predominantly against the hemagglutinin (HA) head (Angeletti and Yewdell 2018). However, the head is highly variable due to antigenic drift, resulting in strain-specific immunity. An improved vaccine should elicit broadly neutralizing antibodies (bnAbs) against conserved sites which not only protect against drifted strains but also various subtypes, preferentially belonging to both group 1 and 2.

In recent years, several attempts have been made to redirect the immune response against the conserved HA stem region, following concepts from reverse and structural vaccinology. Starting from structural information of antibody-antigen complexes, reverse vaccinology proposes the structure-based design of novel vaccine candidates to elicit neutralizing antibodies that are the only known correlates of protection (Burton 2002; Rappuoli et al. 2016, 201; Plotkin 2010). With progress in experimental techniques to isolate and analyze B cells as well as structural characterization of isolated antibodies and their cognate antigens, valuable insights have been gained to facilitate structure-based immunogen design. The reverse vaccinology approach has recently been successfully applied to design novel epitope-focused immunogens for RSV. The obtained results demonstrated that the immune response could be shifted towards subdominant antigenic sites, eliciting neutralizing Abs specific for the targeted epitopes in mice and non-human primates (Correia et al. 2014; Sesterhenn et al. 2019; 2020).

For influenza, several HA stem-targeting bnAbs that are cross-reactive to group 1 and 2 HAs, have been identified (Corti et al. 2011; Dreyfus et al. 2012; Nakamura et al. 2013; Kallewaard et al. 2016; Wu et al. 2015). These antibodies recognize a conserved site on the HA stem domain around the hydrophobic pocket, engaging overlapping epitopes in different orientations. So far, immunogen design approaches for influenza have been mainly focused on removing the immunodominant HA head and stabilizing the stem domain in isolation to overcome the immunodominance of the head. Two promising candidates were reported previously (Impagliazzo et al. 2015) (Yassine et al. 2015) demonstrating that antibodies can be elicited against subdominant epitopes on the HA stem. While these headless designs showed great promise in mounting a broader immune response compared to immunizations with full-length HA, the breadth has mainly been limited to subtypes within groups but low cross-group reactivity was observed (Yassine et al. 2015; Impagliazzo et al. 2015; Steel et al. 2010; Wohlbold et al. 2015; Darricarrère et al. 2021). However, another recent approach, utilizing antigen-reorientation, was more successful at inducing cross-group antibodies (Xu et al. 2024). Here, we investigate whether we could apply structure-based design to engineer mimetics of the conserved HA-stem epitope that can be recognized by a broad panel of bnAbs and elicit cross-reactive immunity in mice. The reported immunogens were designed to capture the surface features of the conserved epitope, only partially relying on structural transplantation of epitope segments while the remaining antigenic surface is mimicked through surface-centric design (Scheck et al. 2022). From the engineered epitope mimetics, two lead candidates were identified that resembled the main properties of the conserved site. We demonstrate that these mimetics can bind to a broad panel of broadly neutralizing, stem-specific antibodies and elicit cross-group antibody response in mice, recognizing HA expressed on virions and on the surface of infected cells. The elicited antibodies are highly specific towards the mimicked site and are cross-reactive to heterologous H1 and H3 strains. Our results show that our surface-centric design approach can be used to engineer epitope mimetics based on heterologous protein scaffolds with precise molecular features of the target epitopes. Furthermore, the epitope mimetics induce broad responses that react with very divergent viral strains and mediating *in vitro* effector functions, showing promise for the use of this strategy in vaccine design challenges that require focusing of antibody responses against conserved epitopes.

## Results

### Design of novel immunogens mimicking a conserved epitope in the hemagglutinin stem

To attempt the design of immunogens that elicit a broad and protective immune response, we selected a well-characterized conserved epitope centered around the hydrophobic pocket on the HA stem. This stem epitope is commonly targeted by bnAbs such as FI6 (Corti et al. 2011), CR9114 (Dreyfus et al. 2012), 39.29 (Nakamura et al. 2013), or MEDI-8852 (Kallewaard et al. 2016) (Figure 1A). The conserved site on the HA stem is a multi-segment epitope, consisting of a 20-residue long α-helix, a four residue VDGW-loop, and a three residue HSV-loop (Figure 1B). We extracted the stem-epitope from a crystal structure of H1 HA in complex with the FI6 Ab (PDB ID: 3ZTN (Corti et al. 2011)) and identified potential protein scaffolds by querying the Protein Data Bank (PDB) (Berman et al. 2000) for proteins with structurally similar backbone segments which were also compatible with the FI6 binding mode (Silva, Correia, and Procko 2016, 201; Correia et al. 2010). However, due to the irregular and discontinuous nature of the epitope, close matches were absent. Based on subsequent searches, we selected a scaffold (PDB ID: 4IYJ (Joint Center for Structural Genomics (JCSG) 2017)) that closely mimicked the α-helix and the VDGW-loop (combined backbone RMSD 1.44 Å), omitting the shorter epitope loop. We hypothesized that the design could mimic core features of the epitope even though close structural matches of the full epitope segments were unavailable. Suitability of the scaffold was confirmed by evaluating the structural compatibility and the predicted binding energy of the scaffold and the FI6 Ab. After transplantation of the epitope helix and VDGW-loop onto the selected protein scaffold, computational sequence design was performed to optimize binding towards the FI6 antibody, resulting in the FI6-focused_01 design (Figure 1C). Initial binding affinity of the computationally designed protein (FI6-focused_01) to FI6 was low, however, specific to the mimicked site when compared to WT protein (Figure S1). To improve binding affinity of the FI6-focused design, we screened a computationally derived combinatorial library including residues adjacent to the grafted epitope helix (Figure S2) for tighter binding to the FI6 antibody and could improve binding to a dissociation constant (K_D_) of 320 nM (FI6-focused_02, Figure S1) measured by Surface Plasmon Resonance (SPR). In the next step, we sought to further increase binding affinity with a site-saturation mutagenesis (SSM) library by sampling positions in the surrounding of the grafted site (aa 93-106 and aa 123-189) that could be relevant for the accurate presentation of the stem epitope (Figure 1C and S2).

**Figure 1.**
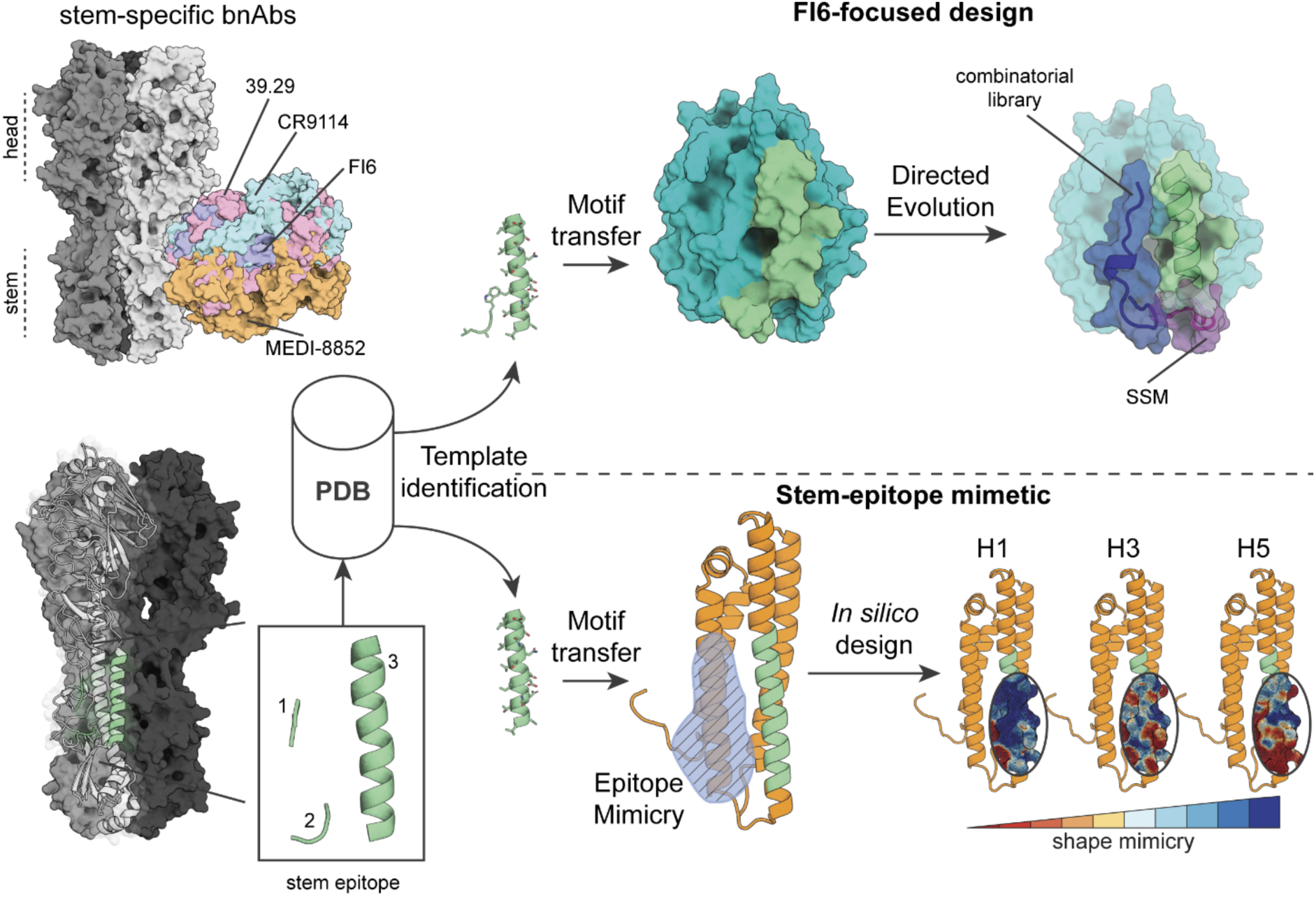
Computational design of stem-epitope immunogens. **Left panel top:** Structures of several bnAbs targeting the conserved site on the immunosubdominant stem of group 1 and 2 HA subtypes. **Left panel bottom:** Stem epitope structural elements. The epitope consists of a short HSV-loop, a VDGW-loop, and a regular α-helix. **Right panel top:** Computational and experimental stages of the FI6-focused design procedure. The helix and the VDGW-loop were queried against the Protein Data Bank (PDB) to retrieve putative scaffolds. The motif was grafted onto a suitable scaffold (PDB ID: 4IYJ) and further improved with directed evolution, both through combinatorial and SSM libraries. Several mutations in the hydrophobic pocket (blue) and a loop connecting the epitope helix to the scaffold (purple) that improved binding affinity to FI6 were revealed by the libraries. **Right panel bottom:** Computational surface-centric design strategy for epitope mimicry. In this design strategy we used only the α-helix for the search of potential protein scaffolds due to the structural complexity of the full epitope. The remaining antigenic site was designed in silico to increase the epitope mimicry and the final design was compared to H1, H3, and H5, demonstrating high epitope mimicry for H1 and H3, and moderate mimicry when compared to H5.

The SSM library approach allows a thorough sampling of a large number of relevant positions. The best individual mutations from this library were combined and we screened 16 variants for improved binding to FI6. All tested variants boosted affinity and the best design, FI6-focused_03, showed a K_D_ of 260 pM, comparable to that of FI6 to H1 HA (Corti et al. 2011) (Figure S1). A second round of computational sequence design was performed, introducing point mutations to monomerize and improve stability of the scaffold, which forms a homodimer in its native state. The final design, FI6-focused_04, was well-folded and monomeric as confirmed by circular dichroism (CD) and multi angle light scattering (MALS) (Figure S3) and bound FI6 with a K_D_ of 6 nM as measured by SPR (Figure 2A). Upon fusion to ferritin nanoparticles it retained FI6 binding (Figure 2B) and analysis by negative staining electron microscopy showed correct assembly of the nanoparticles (Figure 2C).

Following the results of the FI6-focused_04 immunogen design, we concluded that while protein scaffolds closely matching the epitope’s structure were rare, immunogens could be engineered by grafting core epitope elements and improving overall mimicry of the antigenic site through structure-guided design. Accordingly, we focused on the regular epitope α-helix as anchor for new designs, performing scaffold searches solely with the helical segment. As expected, the number of putative scaffolds increased. To identify optimal candidate scaffolds that could maximize epitope mimicry, we applied several filters including predicted binding energy, number of atomic clashes, and number of potential side chain interactions between the scaffold and the FI6 antibody. We identified mouse apolipoprotein E (ApoE) (PDB ID: 1YA9 (Hatters, Peters-Libeu, and Weisgraber 2005)), a regular four-helix bundle that closely mimicked the epitope helix (backbone RMSD 0.5 Å) and offered sufficient surface area to mimic the complete stem epitope (Figure 1D). Since ApoE is involved in lipid transport and highly abundant in serum (Lusis et al. 1987), we hypothesized that the immune response towards the scaffold itself would be low in mice. After grafting the epitope helix side chains onto the scaffold, we evaluated epitope mimicry based on the overall surface mimicry between the protein scaffold and HA with RosettaSurf, and confirmed high similarity to the native epitope (Figure 1D). Based on the observed similarities, computational sequence design was performed to further improve epitope mimicry by 130% over the WT scaffold based on epitope surface shape similarity, resulting in the stem-mimetic_01 design. In addition, epitope mimicry was evaluated by predicting binding energies to CR9114 and MEDI-8852, showing similar values to those of FI6, indicating that the stem-mimetic could engage a range of bnAbs. The stem-mimetic_01 was confirmed to adopt an α-helical fold by CD, was monomeric, and bound with a K_D_ of 44 nM to FI6 (Figure 2D, Figure S3).

To increase immunogenicity and B-cell cross-linking of the FI6-focused_04 and the stem-mimetic_01, the scaffolds were displayed on ferritin nanoparticles (Figure S4). Ferritin assembles from 24 subunits, allowing the multivalent display of proteins resulting in enhanced binding kinetics through avidity (Kanekiyo et al. 2013). While the FI6-focused_04 particle only bound with high affinity to FI6, the stem-mimetic_01 particle showed strong binding to FI6, MEDI-8852 and CR9114, demonstrating its improved binding breadth (Figure 2B, Figure 2E). For both designs well-formed particles could be observed (Figure 2C, Figure 2F). To further enhance the immune response, we fused a known T cell epitope from the influenza matrix protein (M2e) to the N-terminus of the particulate designs (Eliasson et al. 2018) (Figure S4). The production of high-affinity class switched antibodies, as well as the rapid reactivation of memory B cells, requires T cell help (Kurosaki, Kometani, and Ise 2015), however, neither the grafted HA helix nor the ferritin particle is known to provide efficient T cell help (Kelly et al. 2020; Lu et al. 2017).

**Figure 2.**
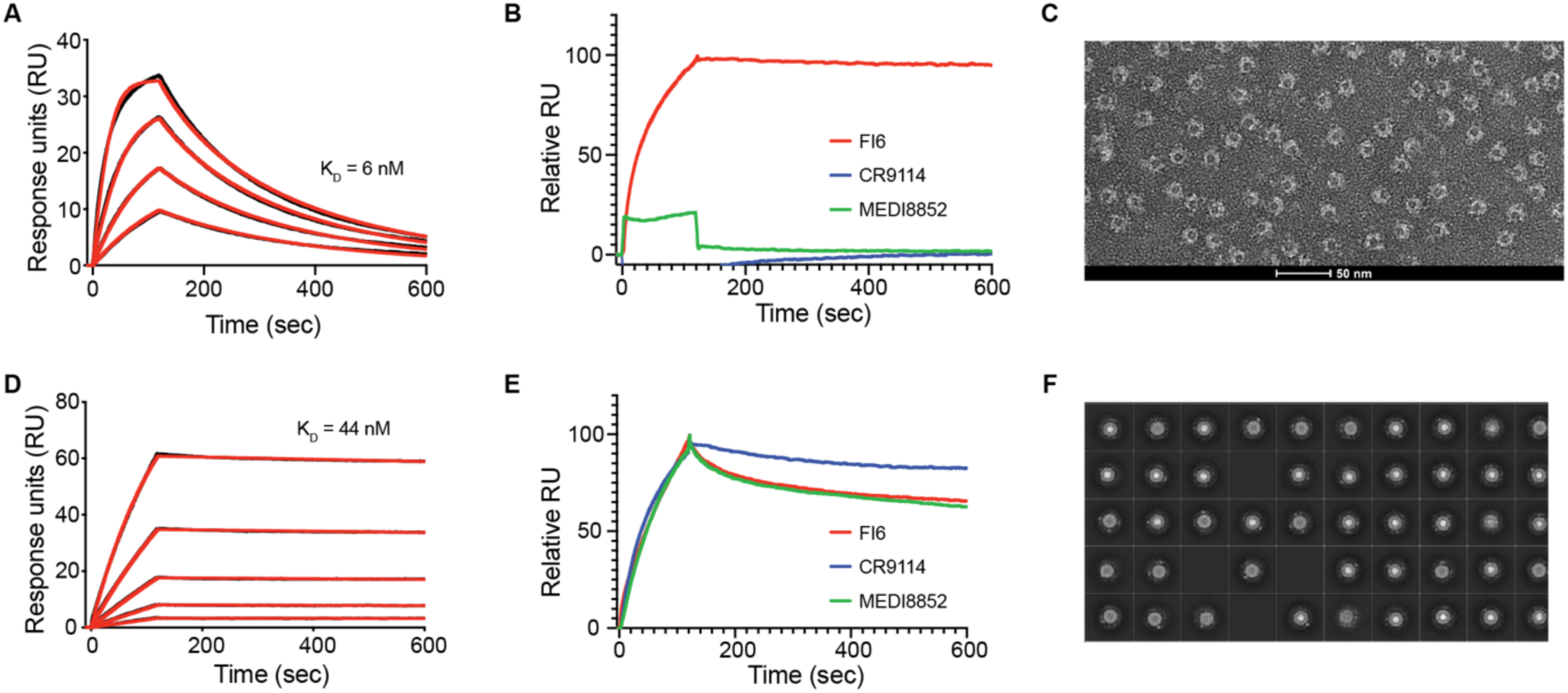
Characterization of designed immunogens. **A.** Surface plasmon resonance (SPR) measurements of the FI6-focused_04 design to FI6 Fab. Analyte concentrations ranged from 62.5 nM to 7.8nM. **B.** SPR measurements for interactions of FI6-focused_04 design, displayed on ferritin nanoparticles, with stem-specific antibodies, FI6 (red), CR9114 (blue) and MEDI-8852 (green). **C.** Negative stain electron (EM) of FI6-focused_04 design on ferritin nanoparticles showed assembly of the nanoparticle and presentation of the design. **D.** SPR measurements of the stem-mimetic_01 design to FI6 Fab. Analyte concentrations ranged from 5 µM to 312.5 nM. **E.** SPR measurements for interactions of stem-mimetic_01 design, displayed on ferritin nanoparticles, with stem-specific antibodies, FI6 (red), CR9114 (blue) and MEDI-8852 (green). **F.** Negative stain EM of the stem-mimetic_01 on ferritin nanoparticles showed correct assembly of the nanoparticle and presentation of the design.

### Structural characterization of designed stem-epitope mimetics

To visualize the structure of the designed immunogens and their binding to stem-antibodies, we solved the crystal structure of the FI6-focused_03 design in complex with FI6 Fab at 1.95 Å resolution (Figure 3A). Comparing the solved structure and designed model demonstrated close structural similarity with a RMSD of 2 Å. We observed that the grafted epitope helix was elongated by two additional turns at the N-terminal end. This is likely attributed to a mutation introduced as part of the epitope (P144T), forming the transition of helix to loop in the native scaffold. However, the introduced mutation benefits the interaction with FI6 and mimicry of the stem epitope was still high when compared to H1 HA (Figure 3B).

In parallel, we solved the crystal structure of stem-mimetic_01 in complex with CR9114 to a resolution of 2.7 Å (Figure 3C). Comparison of structure and model revealed close structural agreement with a RMSD of 1.6 Å. Close inspection of the side chain placement revealed that rotamers generally adapted the designed conformations and many of the polar and hydrophobic interactions were accurately retained. We compared the structure of our design to H5 in complex with CR9114 (PDB ID: 4FQI (Dreyfus et al. 2012)). Polar interactions of the residues surrounding the epitope helix were partially recovered, while polar interactions formed by the epitope helix were conserved (Figure 3D). Additionally, the epitope surface of our stem-mimetic’s crystal structure resembled the surface of H5 HA complex with CR9114 (Figure 3E), demonstrating successful mimicry of the epitope.

**Figure 3.**
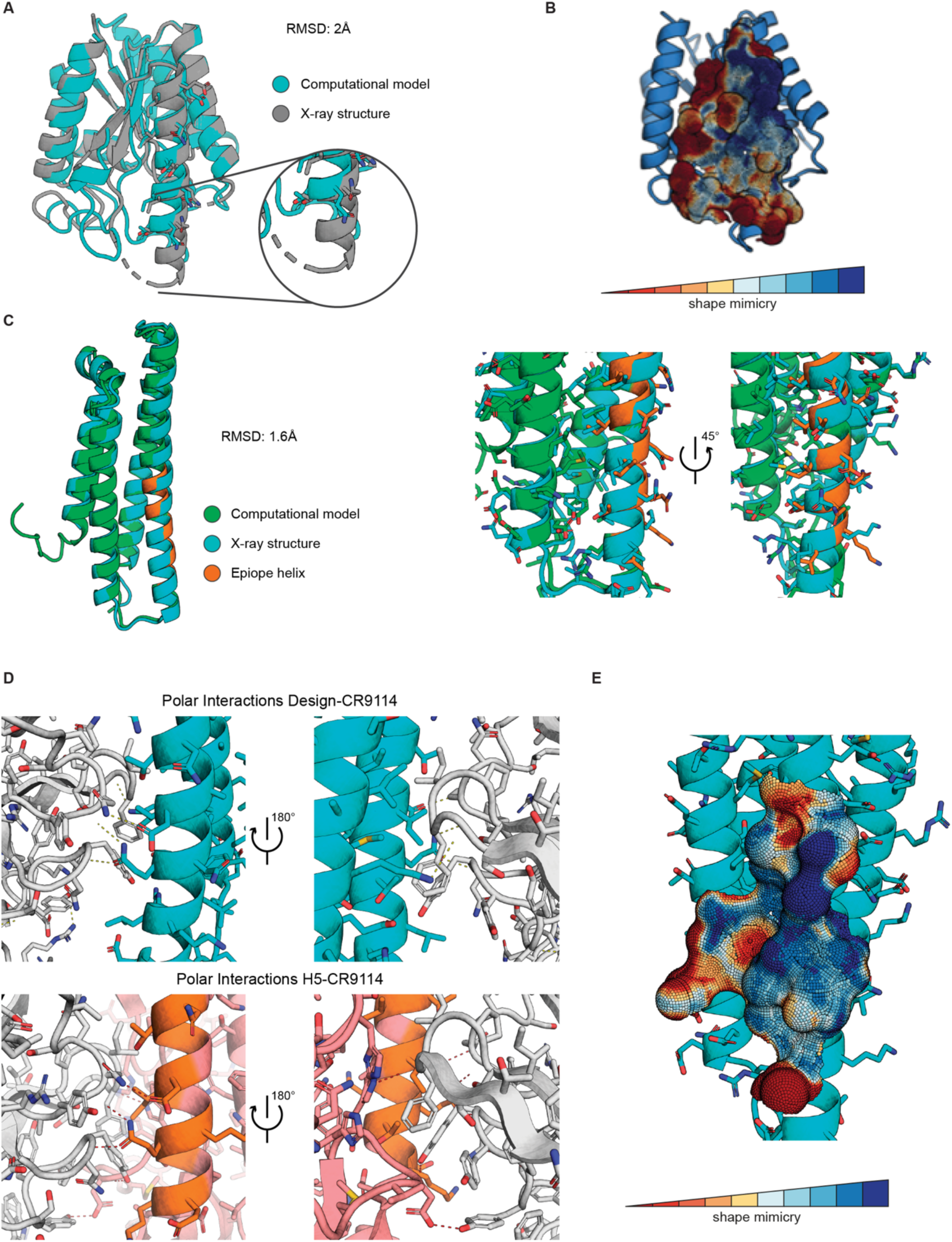
Structural characterization of designed immunogens. **A.** Structural comparison of FI6-focused_03 model to X-ray structure shows overall agreement with a RMSD of 2 Å, however, two additional helix-turns were formed on the epitope-helix N-terminal end. **B.** High epitope mimicry of X-ray structure and H1. **C.** Structural comparison of the stem-mimetic_01 computational model to the solved X-ray structure in complex with CR9114. The computational model showed excellent structural agreement with the solved X-ray structure (RMSD 1.6 Å), confirming the recovery of modeled rotamer positions. **D.** Polar interactions of the stem-mimetic in complex with CR9114 were compared to a H5-CR9114 complex, showing full recovery of interactions involving the epitope helix and partial recovery for the remaining epitope. **E.** The epitope region was well mimicked in the structure when comparing surface shape similarity to the corresponding H5 stem epitope.

### Selective pull-down of cross-reactive antibodies from human PBMCs

To assess the capability of both designs to engage known anti-HA antibody lineages, we investigated their ability to bind memory B cells in human peripheral blood mononuclear cells (PBMCs). We used samples from one individual enrolled in the VRC 310 clinical trial, which received H5 DNA-prime/monovalent inactivated H5N1 boost vaccination (Ledgerwood et al. 2011; 2013). For the FI6-focused_03 design, memory B cells cross-reactive to the design and an H1 stem construct (Andrews et al. 2023) were sorted (Figure 4A). For the stem-mimetic, design positive memory B cells were sorted in a H1 only and a H1/H3 double positive population and sequenced (Figure 4B).

A major population of pulled down antibodies from both designs used the VH3-23, a germline gene that is known to give rise to potent bnAbs (Joyce et al. 2016). All B cells in this population were clonally related. A representative antibody of this clone (31-1B01) had a remarkable binding breadth, similar to FI6 (Figure S5B). The FI6-focused design also pulled down antibodies utilizing the VH3-30 gene, similar to FI6 (Figure 4A). However, these antibodies were, in contrast to FI6, mainly restricted to group 1 HAs (Figure S5B). Sorting of B cells from a second donor from the same clinical trial showed an even stronger focusing to VH3-30 antibodies (Figure S5A), which were also mainly specific for group 1 HAs. The pan-group binding of FI6 is the result of affinity maturation caused by somatic mutations, germline FI6 (FI6-GL) is restricted to group 1 (Corti et al. 2011), in line with our observations that all sorted antibodies from VH3-30 were binding predominantly to group 1 HAs (Figure S5B).

While all of the stem-mimetic positive B cells that were cross-reactive to H1 and H3 were from the same lineage (represented by 31-1B01), the H1 only population mainly contained antibodies from VH1-69 (Figure 4B), which represents the major human VH region giving rise to group 1 specific HA stem antibodies. The stem-mimetic’s ability to engage this class of antibodies could be beneficial for the robust induction of pan-group 1 bnAbs.

Even though binding to memory B cells is not a guarantee for effective activation and induction of neutralizing antibodies, our results show that our design can engage relevant human antibody lineages that can provide broad influenza neutralization.

**Figure 4.**
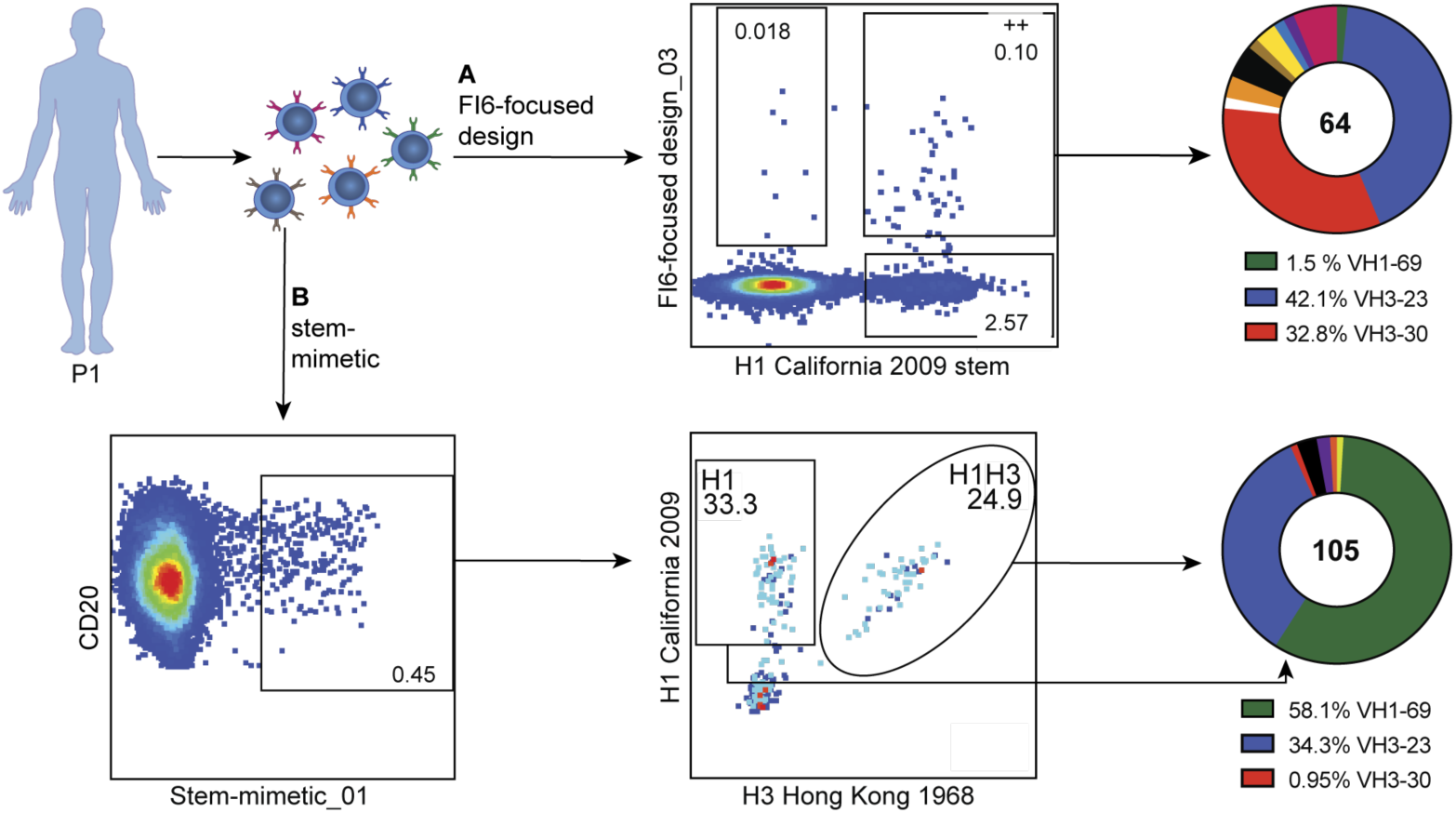
Antibody pull down from human PBMCs. **A.** Memory B cells cross-reactive to the FI6-focused design and H1 were sorted and sequenced and their VH sequences were assigned to their originating germline regions for comparison with known bnAbs. **B.** Stem-mimetic positive memory B cells were sorted in a H1 only and a H1/H3 double positive population and both populations were sequenced to assign their corresponding VH genes.

### Elicitation of stem-specific antibodies using computationally designed stem epitope antigens

To investigate the ability of our designs to elicit antibodies *in vivo*, naïve BALB/c mice were injected three times with nanoparticles displaying either FI6-focused_04 or stem-mimetic_01 immunogen, adjuvanted with AS03 (Figure 5A) (Garçon 2012). Both designs were immunogenic as seen by the high design-induced antibody titers. Since our designs were based on naturally occurring, heterologous proteins, we used the respective WT scaffolds to determine the proportion of antibodies targeting the epitope. The antibody response against the WT scaffolds reflects the proportion of non-epitope-specific antibodies. Thus, a lower response to the WT scaffold correlates with high epitope-focusing. Testing antibody titers against the WT scaffold of the stem-mimetic_01 immunogen confirmed strong epitope-focusing (Figure 5B). As the scaffold protein of stem-epitope_01 is an endogenous mouse protein (Hatters, Peters-Libeu, and Weisgraber 2005) we hypothesized that elicited antibodies were epitope-specific and did not target the scaffold due to B-cell tolerance. The immuno-silent scaffold protein renders this design an ideal immunogen, as it promotes a commonly subdominant epitope to become immunodominant.

In the next step, the elicited antibodies from both designs were tested for reactivity to different full-length HAs from both phylogenetic groups. Both designs showed comparable titers to H1, however, stem-mimetic_01 immunized animals showed significantly higher titers to H3 (Figure 5C). Nevertheless, in the same animals, the titers against H3 are considerably lower than against the stem-mimetic_01 itself, even though the majority of antibodies were specific for the grafted epitope (Figure 5B). This discrepancy is most likely resulting from antibodies which approach the epitope in an angle that is not accessible on full-length HA, similar to results from previous studies (Schneemann et al. 2012). To determine the epitope focusing on full-length HA, we tested the sera against a mutated version of H1 (H1Δstem) containing a glycan in the stem epitope which has been reported to disrupt binding by stem-specific Abs (Darricarrère et al. 2021). For both designs the glycan knocked out binding almost completely, confirming the specificity of the elicited antibodies to the selected stem epitope (Figure 5D). In contrast, sera from mice immunized with three times HA full-length (Methods and Materials) did not show decreased binding to H1Δstem, demonstrating the poor response against this subdominant site. The results are supported by earlier studies reporting that even after immunization with headless HAs, the stem epitope is targeted poorly in mice (Sangesland et al. 2019). Due to the distinct epitope focusing and the close mimicking of H3 we decided to concentrate further experiments on the stem-mimetic_01.

To further analyze the induced humoral response on a cellular level, we isolated B cells from mice immunized with stem-mimetic_01 particles and evaluated their cross-reactivity to an H1 and an H3 HA. Spleens and draining lymph nodes were examined two weeks after the 2^nd^ and the 3^rd^ injection and class switched B cells were analyzed for design and HA binding (Figure 5E and Figure S6). While the proportion of design positive B cells remained stable between the 2^nd^ and 3^rd^ injection, a clear increase of the H1/H3 cross-reactive antibodies after the 3^rd^ injection could be observed. The results furthermore highlight that the induced HA positive antibodies are almost entirely cross-reactive to H1 and H3, where the induction of such bnAbs has been historically difficult in mice and could not be achieved with H1 and H3 mosaic nanoparticles (Cohen et al. 2021).

**Figure 5.**
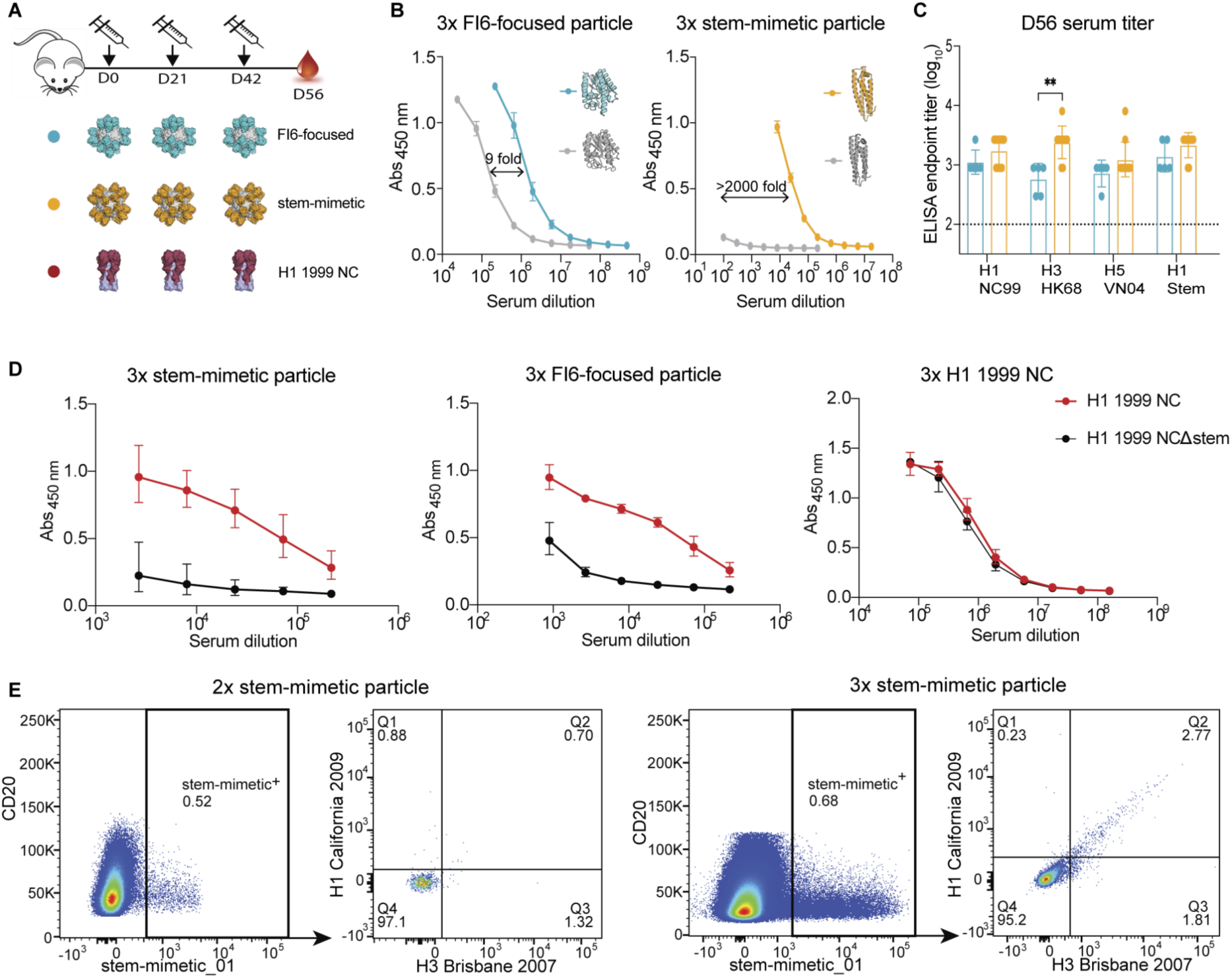
Design induced antibody responses. **A**. Immunization scheme and immunogens used in this study (blue = FI6-focused_04, orange = stem-mimetic_01). FI6-focused_04 and stem-mimetic_01 immunogens were presented on ferritin nanoparticles, H1 HA was given as soluble trimers. All immunogens were adjuvanted with AS03 and injected intramuscularly three times in three week intervals. **B.** Antibody titers measured against the respective design (blue or orange) and WT scaffold (grey) in sera from final bleeding (day 56 after first injection). Plates were coated with monomeric designs. The mean values and SD (n=5) are shown. **C.** Cross-reactive titers against HA. Sera from D56 tested against H1 1999 NC, H3 1968 HK, H5 2004 VN and H1 1999 NC stem HAs. Plates were coated with trimeric HAs. The mean values and SD (n=5) are shown. ** p<0.01. **D.** Antibody titers against H3 1968 HK and pH1 2009 CA UV-inactivated virus. The mean values and SD (n=5) are shown. **E.** Epitope focusing on H1 HA. Binding curves against H1 1999 NC and H1 1999 NC stem KO mutant. Plates were coated with trimeric HAs **F.** Cross-reactivity to H1 and H3 of stem-mimetic positive mouse B cells in the draining lymph nodes after 2 and 3 immunizations with stem-mimetic_01 particle.

### Induction of cross-reactive responses in mice using stem-mimetic antigen

To examine if the elicited antibodies not only bind to recombinant antigens but also to native antigens on virions, whole-cell virus ELISA was performed. Mice were immunized in a homologous boost scheme with stem-mimetic nanoparticle or WT HA (H1 or H3), or a heterologous prime boost immunizations with H1 WT trimers as prime injections and stem-mimetic nanoparticle or PBS as boost injections. A final heterologous boost scheme was performed with a first injection of stem-mimetic particle and HA (H1 or H3) boost injections (Figure 6A). ELISA plates were coated with UV-inactivated virus and binding titers were determined. Serum antibodies elicited by stem-mimetic_01 showed virus binding on par with three times H1 full-length trimer immunization against a heterologous pandemic H1 (A/California/07/2009) virus and a more recent H1 strain (A/Michigan/45/2015) (Figure 6B). Notably, priming with stem-mimetic_01 and H3 boost induced antibodies exhibiting superior binding to an H3 HK68 and H3 Singapore (A/Singapore/INFIMH-16-0019/2016) viruses (Figure 6B). Binding was also observed to an avian H7 virus (A/Environment/Suzhou/SZ19/2014) not circulating in the human population, highlighting the potential breadth the stem-mimetic particle induces (Figure 6B). Similar results were obtained using a flow cytometric assay to detect binding to nascent HA on infected MDCK cells (Figure 6C, Fig. S7A-B), showing a higher degree of binding after a heterologous boost with the stem-mimetic particle. Considering that the stem-epitope mimetic carries a single antigenic site representing only ∼6% of the HA surface area, the enhanced binding breadth of the elicited antibodies is remarkable.

Stem-specific antibodies are known to be weakly neutralizing and mainly protect via Fc-receptor mediated cellular pathways (DiLillo et al. 2016). Therefore, we evaluated the activation of antibody-dependent cellular cytotoxicity (ADCC) by the elicited antibodies in mouse sera. We compared homologous immunizations with stem-mimetic nanoparticle *vs* heterologous prime-boost with WT H1 or H3 as prime and boosted with stem-mimetic nanoparticle or PBS (Figure 6A). HA prime only injections did not result in antibodies with ADCC activity against any of the three tested viruses, H1N1 PR8, H1N1 Ca07/09 and H3N2 X31, while sera from all groups boosted with the stem-mimetic particle showed ADCC activity against all viruses (Figure 6D). Furthermore, stem-mimetic only immunization was able to induce ADCC-activating antibodies in the majority of mice tested (Figure 6D). The results show that the stem-mimetic is able to induce functional cross-reactive antibodies against H1 and H3 viruses and boost pre-existing HA antibodies in a heterologous prime boost scheme. Strikingly, the heterologous prime boost injections showed more consistent ADCC activation. Remarkably, 3x stem-mimetic particle injections resulted in robust ADCC activity against both H1 viruses and partially against the H3 virus, highlighting the potency of the stem-mimetic design alone.

**Figure 6.**
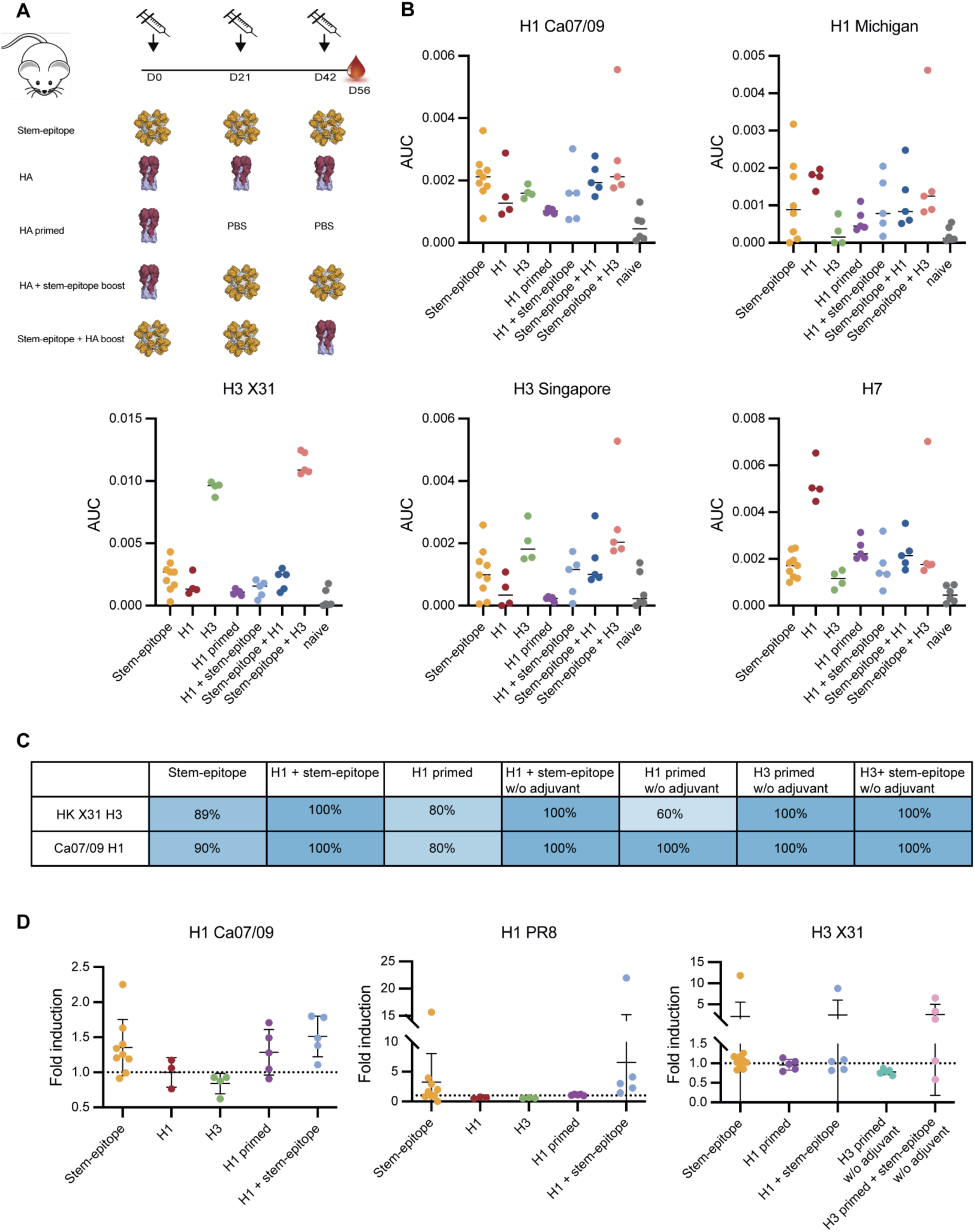
Response breadth mediated by stem-mimetic. A. Homologous immunizations were performed with stem-epitope particle, H1 (NC99) trimer, or H3 (HK68) trimer. Prime injections were performed with H1 (NC99) trimer or H3 (HK68) trimer and boosted with PBS. Boost injection schemes were performed with a H1 (NC99) trimer or H3 (HK68) trimer prime and stem-mimetic particle boost. HA boost injection schemes were performed with two immunizations of stem-epitope particle and a third immunization with H1 (NC99) trimer or H3 (HK68) trimer. **B.** Antibody titers against H1 1999 NC, H1 2015 Michigan, H3 X31, H3 2016 Singapore and H7 2013 Shanghai UV-inactivated virus. Median values are represented by horizontal bar. **C**. Percentage of mice per group producing sera that bind to virus infected MDCK cells. **D.** Antibody-dependent cellular cytotoxicity (ADCC) mediated by design-induced antibodies against three different influenza viruses. Relative luminescence units (RLU) were determined for all groups and the fold induction over background (negative control) was calculated. The mean values + SEM for all mice in the different groups are shown (n = 5, except 3 x stem-epitope particle n = 10).

## Discussion

We have developed novel stem-epitope mimetics that elicit stem-specific antibody responses against group 1 and group 2 influenza subtypes. Our work demonstrates that computational protein design centered around mimicking antigenic surfaces offers a new approach to present structurally complex epitopes that are irregular and consist of multiple segments. Computational design resulted in two immunogens with different binding breadths to various stem-targeting antibodies. Our FI6-focused design was based on the structural modeling of the FI6 epitope helix and VDGW-loop, while we attempted to recover the remaining epitope area through additional design steps. A previous study using only the epitope helix failed to bind relevant antibodies or to induce protective antibodies in mice (Schneemann et al. 2012). The predicted structure of the immunogens was confirmed by X-ray crystallography, demonstrating close structural resemblance of the crystallized structure and model. The initial design bound with low affinity to the FI6 antibody but interactions were improved through directed evolution to obtain a low nanomolar binder. As a result of the strong selection bias towards the FI6 antibody, the immunogen was highly-specific to FI6 but did not readily interact with other known stem-specific antibodies.

Since appropriate structural templates are rare for this type of multi-segment epitope (Bonet et al. 2018), we further focused on transplanting only singular, regular structural elements while emphasizing computational design steps to mimic the entire antigenic site through surface design. To this extent, we identified a mouse apolipoprotein E that forms a four-helical bundle, ideal for fitting the epitope α-helix in a stable and correctly oriented conformation. The scaffold offered a sufficiently large additional contact area to mimic central parts of the complete antigenic site. Rational design was performed to improve the similarity of our design and the stem epitope. Remarkably, computational design was sufficient to attain an immunogen that not only bound to FI6 with high affinity but also engaged additional stem-specific antibodies like CR9114 and MEDI-8852. X-ray crystallization of the designed immunogen showed excellent structural agreement with the computational model. The structure was obtained in the presence of CR9114 and was shown to recover most polar and hydrophobic interactions that were observed in a native H5-CR9114 complex.

The large majority of the population has been exposed to influenza at least once in their life, thus possessing pre-existing immunity. Studies have shown that broadly neutralizing antibodies can routinely be isolated from recently infected or vaccinated donors (Joyce et al. 2016; Andrews et al. 2017). However, since these antibodies occur at low abundance they do not confer protection against varying influenza viruses. Even though individuals produce memory B cells, they are not sufficiently expanded in secondary responses (Andrews et al. 2015; H. Tan et al. 2019; Guthmiller et al. 2020). One of the main reasons for this outcome is likely related to the immunodominance of the HA head domain. While most of the bnAbs are targeting a conserved epitope on the stem domain of HA, humoral immune responses are highly skewed towards the variable head domain (Angeletti et al. 2017). Therefore, enhancing pre-existing stem-specific bnAbs could potentially increase their abundance and confer broad protection to varying influenza strains. We demonstrated that both designed immunogens are able to bind to pre-existing bnAbs in human PBMCs, corroborating their potential to reactivate existing B cells. While the FI6-focused design showed a high specificity for VH3-30 antibodies, the stem-mimetic was mainly bound by VH1-69 antibodies. Since this VH region encodes for the majority of group 1 specific bnAbs targeting the HA stem domain in humans, being able to selectively engage and boost this antibody class could provide a potent antibody response against group 1 influenza viruses. Nevertheless, using the stem-mimetic, we were able to pull down a potent mAb, 31-1B01, which makes use of the VH3-23 gene and was able to bind all tested HA.

Mouse immunizations showed that both designs induced antibodies cross-reactive to H1 and H3 HAs. In general, the observed antibody titers after immunization with designed constructs were lower than native HA induced titers. This is to be expected since the immunogens represent a single antigenic site on HA that is subdominant while full-length HA hosts additional epitopes. Additionally, the limited ability of mice to produce antibodies against the targeted epitope has been demonstrated before (Sangesland et al. 2019). The VH gene giving rise to the majority of bnAbs against the HA stem is VH1-69 (Andrews et al. 2017; Joyce et al. 2016), which according to our antibody pull down results is engaged by our stem-mimetic. However, mice do not carry this germline and are therefore not able to induce this class of antibodies.

Until now only a few mouse antibodies were found targeting this epitope and none of them recognized both group 1 and 2 HAs (G. S. Tan et al. 2014; Hai et al. 2012; Okuno et al. 1993). Further, pan-group mAbs are also exceedingly rare in humans (Joyce et al. 2016). However, binding of mouse B cells to H1 and H3 probes after stem-mimetic immunizations and other *in vitro* assays demonstrate that design-induced antibodies are able to cross-react with group 1 and 2 HAs present on virus particles and infected MDCK cells. Although our stem-mimetic only displays a single epitope from HA and did not confer protection from a lethal challenge (data not shown), it is able to induce H1-H3 cross-reactive ADCC responses. These results reinforce the potential of our stem-mimetic to elicit a highly focused and functional antibody response, superior to immunizations with native antigens.

To increase robustness of protection, we hypothesize that combining several distinct antigenic sites into a single formulation may lead to an improved immune response. For respiratory syncytial virus (RSV) it has been shown that an immunization strategy with three designed immunogens displaying three distinct epitopes of the RSV F glycoprotein induces neutralizing antibodies in mice and non-human primates (Sesterhenn et al. 2020). The cocktail formulation of the immunogens was shown to elicit an epitope-focused immune response and was able to reshape the antibody response when used as a boosting agent. Combining non-overlapping epitopes from the HA protein, covering the head and stem region could potentially induce similar immune responses. Furthermore, our work highlights that computational design allows the mimicry of epitopes even if close structural resemblance of the entirety of the epitope is absent but the resulting surfaces are similar. This approach is not only limited to the design of immunogens but can be applied more generally to the design of functional proteins, especially for irregular and discontinuous structural motifs.

## Acknowledgments

We thank Ying Huang, Karen Matsuoka, and Ankita Balsaraf for expression, harvesting, buffer exchange, and purification of CR9114 Fab from culture supernatants and purification of stem-mimetic_01 from *E. coli*. We thank Trissa Elkins and Robert Nolte for crystallography and computing support. We additionally acknowledge EPFL core facilities for negative staining, the GECF for sequencing support, the CPG for mouse immunization support. The computational design was supported by EPFL Scientific IT & Application Support (SCITAS) and the Swiss National Supercomputing Centre (CSCS). The work was supported by the Swiss National Science Foundation. We acknowledge Protein Production Sweden for provisioning of facilities and experimental support, and we would like to thank M. Andersson and M. Bäckström for assistance. Protein Production Sweden is funded by the Swedish Research Council as a national research infrastructure. We thank the staff at the Experimental Biomedicine (EBM) core facility at the University of Gothenburg for animal management. The following viruses were obtained via the Influenza Reagent Resource (IRR), CDC, USA: A/Michigan/45/2015, A/Singapore/INFIMH-16-0019/2016. We thank Qiyun Zhu (Chinese Academy of Agricultural Science) for the UV inactivated A/Environment/Suzhou/SZ19/2014. This research used resources of the Advanced Photon Source; a U.S. Department of Energy (DOE) Office of Science User Facility operated for the DOE Office of Science by Argonne National Laboratory under Contract No. DE-AC02-06CH11357. Data were collected at Southeast Regional Collaborative Access Team (SER-CAT) 22-ID (or 22-BM) beamline at the Advanced Photon Source, Argonne National Laboratory. SER-CAT is supported by its member institutions (see www.ser-cat.org/members.html), and equipment grants (S10_RR25528 and S10_RR028976) from the National Institutes of Health. S.F.A., F.M., and A.M. are supported by the Division of Intramural Research, NIAID, NIH. D.A. is supported by grants from the European Research Council (B-DOMINANCE, grant no. 850638), the Swedish Research Council (grants nos. 2017-01439, 2021-01164 and 2021-01165), the Sahlgrenska Academy and the Institute of Biomedicine at the University of Gothenburg.

## Author contributions

S.W., A.S., and B.E.C. conceived the work and designed the experiments. A.S. performed the computational design and S.W. performed the experimental characterization. L.R. performed the mouse B cell staining and virological experiments under the supervision of D.A. F.M. pulled down and analyzed antibodies from human PBMCs under the supervision of S.F.A. and A.M. J.C. solved crystal structures of the FI6-focused design complex under the supervision of T.K.. W.H. and E.M. solved the crystal structure of the stem-mimetic complex. S.W., S.G., S.R., K.S. and L.R. performed experiments and conducted animal studies. B.B., C.P.M, R.R., N.B. and V.V. contributed to the design and planning of animal studies. S.W., A.S., K.M.C. and B.E.C wrote the manuscript with input from all authors.

## Data availability

Data supporting the findings of this manuscript are available from the corresponding authors upon reasonable request. The atomic coordinates and corresponding structure factors have been deposited to the RCSB Protein Data Bank under PDB ID 8UZP (stem-mimetic_01-CR9114 Fab), 9HI4 (design_03 scaffold in complex with FI6-Fab), 9HI5 (scaffold FI6-focused design_03) and 9HI6 (scaffold FI6-focused design_04 in complex with FI6-Fab).

## Methods

### Computational design of stem-epitope mimetics

#### FI6-focused designs

The structural segments comprising the conserved stem epitope around the hydrophobic pocket were extracted from the H1 crystal structure in complex with the FI6 antibody (PDB ID: 3ZTN). The epitope consists of three segments, a three residue HSV-loop (residues 28-30, chain A), a four residue VDGW-loop (residues 18-21, chain B), and an α-helix (residues 38-57, chain B). We performed a structural search of the epitope against the Protein Data Bank (PDB, version August 2018) containing 141,920 protein structures to identify putative scaffold candidates based on the local similarity. The search was performed using the MASTER software (Zhou and Grigoryan 2015) with a backbone RMSD threshold below 2 Å, however, no suitable scaffolds were detected according to both, local structural features or overall topology. A second search was performed, omitting the HSV-loop of the epitope to increase chances of local structural matches, resulting in 45,616 matches with backbone RMSD below 2 Å. The potential scaffold set was narrowed down by restricting the protein length to 50-250 residues and evaluating the accessibility of the epitope to the FI6 antibody in terms of predicted binding energy and atomic clashes. The remaining candidates were inspected manually to select scaffolds which present the epitope in its native conformation and provide additional surface area to mimic the entire antigenic site. We selected a putative acylhydrolase (PDB ID: 4IYJ) that matches to the trimmed epitope with a RMSD of 1.44 Å and transplanted the epitope side chains of the α-helix and VDGW-loop onto the scaffold (FI6-focused_01). We expressed 45 design variants on yeast and screened binding to the FI6 antibody. Based on the screening on the yeast surface, we identified one design that showed specific interaction with the antibody, albeit with low affinity. Since initial binding of the design protein to the FI6 antibody was low, we improved binding through a combinatorial library by sampling positions adjacent to the epitope helix, resulting in a 320 nM K_D_ binder (FI6-focused_02). The binding affinity of the FI6-focused design was further improved by a subsequent SSM library, sampling epitope positions and surrounding residues (aa 93-106 and aa 123-189). We combined best individual mutations to improve binding affinity with the least amount of additional mutations. In total 16 variants were screened for improved binding to FI6 and all design variants boosted affinity, with the best binding design, FI6-focused_03, resulting in a K_D_ of 0.26 nM. Since the native protein scaffold forms a homodimer, we introduced several mutations to disrupt dimer formation. Monomerization was necessary to enable efficient display on protein nanoparticles which will be used for immunization studies. The native homodimerization interface of the protein scaffold is located opposite to the transplanted epitope. Residues contributing to the dimerization (ddG < -0.8) and exposed hydrophobic residues in the interface were selected and submitted to sequence design. We generated 50 models and selected the best design based on overall Rosetta energy units (REU). We selected eight mutations that drastically decreased the predicted binding energy, suggesting disruptive interactions, while maintaining an overall energy for the model on par with the native monomeric subunit. To improve stability of the resulting monomeric design variant, we performed a BLAST search (NCBI Resource Coordinators 2018) of the WT protein scaffold sequence and constructed a position-specific scoring matrix (PSSM) used during subsequent sequence design. We selected mutations that improved local residue REU and were not part of the epitope region, resulting in twelve mutations in total to increase thermostability of the protein, resulting in the final design, named FI6-focused_04.

#### Stem-epitope mimetics

Based on previous results that indicated a lack of suitable protein scaffolds for the complex epitope, we decided to focus on identifying scaffolds based on the regular α-helix. We performed a scaffold search against a subset of the PDB containing monomeric proteins with helical SSEs and a length of 50 to 250 amino acids with a RMSD cutoff at 1 Å, similar as described above but using Rosetta MotifGraft protocol (Silva, Correia, and Procko 2016) to align the structures, resulting in 7,655 matches. To narrow the set of potential scaffolds we lowered the RMSD threshold to 0.3 Å and allowed no steric clashes, resulting in 1,525 matches. Potential scaffolds were further filtered by assessing accessibility of the FI6 antibody and computing the number of putative contacts between the scaffold and antibody in the epitope region to evaluate the potential to improve overall epitope mimicry. Based on these selection criteria we manually evaluated the top 50 matches and selected a mouse apolipoprotein E (ApoE, PDB ID: 1YA9) as design candidate. The side-chains of the epitope helix were transplanted onto the scaffold using Rosetta MotifGraft and three mutations on the scaffold, not part of the interface, were introduced to resolve steric hindrance with epitope residues. Next, we evaluated the overall epitope mimicry based on surface similarity using RosettaSurf (in preparation) to identify positions with low epitope mimicry. We identified four interface positions that demonstrated low resemblance of the antigenic site and subjected these positions to surface-centric design. We performed RosettaSurf-site on these positions, sampling all 20 amino acids at each position and evaluating their impact on overall epitope mimicry. Individual mutations were evaluated based on their shape similarity to the antigenic site as observed in the H1-FI6 complex (PDB ID: 3ZTN), their propensity to form α-helices (Chou and Fasman 1978), and their biochemical properties, preferring hydrophobic residues to mimic the hydrophobic pocket of the epitope. Designed variants were evaluated based on predicted binding energy to the FI6 antibody, followed by manual inspection of the best ranking designs. A rational mutation I98A was introduced, as the designed amino acids did not match the native epitope residues, resulting in the stem-epitope_01. An additional variant (stem-epitope_02) introducing two potential disulfide bonds connecting positions 53-97 and 96-137, for improved stability was ordered. The proteins were recombinantly expressed in E. coli and purified. Both proteins were monomeric, correctly folded, and bound to antibody FI6 with K_D_s of 44 nM and 48 nM, for the stem-epitope_01 and stem-epitope_02, respectively. The introduced disulfide bonds in stem-epitope_02 did not increase stability as evaluated by its melting temperature and thus stem-epitope_01 was used for further analysis.

Data analysis was performed with the help of the rstoolbox Python library (Bonet et al. 2019) and protein structures were visualized using PyMOL (Schrödinger, LLC 2015).

#### Yeast surface display of single designs

DNA sequences of all designs were purchased from Twist Bioscience containing homology overhangs for cloning. DNA was transformed with linearized pCTcon2 vector (Addgene #41843) into EBY-100 yeast using the Frozen-EZ Yeast Transformation II Kit (Zymo Research). Transformed yeast were passaged once in minimal glucose medium (SDCAA) before induction of surface display in minimal galactose medium (SGCAA) overnight at 30°C. Transformed cells were washed with PBS + 0.05% BSA and incubated with different concentrations of FI6 antibody for 2h at 4°C. Cells were washed once and incubated for additional 30 min with FITC-conjugated anti-cMyc antibody (sigma, #SAB4700448) and PE-conjugated anti-human Fc (invitrogen, #12-4998-82). Cells were washed and analyzed using a Gallios flow cytometer (Beckman Coulter).

#### Combinatorial libraries

Combinatorial sequence libraries were constructed by assembling multiple overlapping primers containing degenerate codons at selected positions for combinatorial sampling of the epitope. Primers were mixed (10 µM each), and assembled in a PCR reaction (55°C annealing for 30 sec, 72°C extension time for 1 min, 25 cycles). To amplify full-length assembled products, a second PCR reaction was performed, with forward and reverse primers specific for the full-length product. The PCR product was desalted and used for transformation.

#### Yeast surface display of libraries

Combinatorial libraries and SSM libraries were transformed as linear DNA fragments in a 5:1 ratio with linearized pCTcon2 vector as described previously (Chao et al. 2006) into EBY-100 yeast. Transformation efficiency generally yielded around 10^7^ transformants. Library cultures were prepared for sorting similar to single designs. Labelled cells were sorted on a Sony SH800 cell sorter. For combinatorial libraries, sorted cells were grown in SDCAA and prepared similarly for two additional rounds of sorting. After the 3^rd^ sort cells were plated on SDCAA plates and single colonies were sequenced. SSM libraries were only sorted once and grown in liquid culture for plasmid prep.

### Protein expression and purification

#### Design scaffolds

DNA sequences were purchased from Twist bioscience and cloned into a pET21b vector for bacterial expression with a c-terminal 6x His Tag. Plasmids were transformed in *E. coli* BL21 (DE3) and grown overnight in LB medium supplemented with Ampicillin. Overnight cultures were used to inoculate the main culture at an OD_600_ of 0.1. Cells were grown at 37°C till they reached an OD_600_ of 0.7 and induced with 1mM IPTG. FI6-focused design versions were incubated for 4-5h at 37°C. Stem-epitope design was incubated overnight at 22°C. Cultures were harvested by centrifugation. Pellets were resuspended in lysis buffer (50 mM Tris, pH 7.5, 500 mM NaCl, 5% Glycerol, 1 mg/ml lysozyme, 1 mM PMSF, and 1 µg/ml DNase) and sonicated on ice for a total of 12 minutes, in intervals of 15 s sonication followed by 45 s pause. The lysates were clarified by centrifugation (48,000 *g*, 20 min) and purified via Ni-NTA affinity chromatography followed by size exclusion on a HiLoad® 16/600 Superdex® 200pg column on an ÄKTA™ pure system (Cytivia).

#### Antibodies

The DNA sequence of all used human antibodies were ordered from Twist Bioscience and cloned into a pHLsec vector for mammalian expression containing a C-terminal human Fc fragment for heavy chain cloning and no Tag for light chain cloning. Antibodies were produced using the Expi293™ expression system from Thermo Fisher Scientific. Supernatant was collected 6 days post transfection and purified via protein A affinity chromatography and subsequent size exclusion on a HiLoad® 16/600 Superdex® 200pg column on an ÄKTA™ pure system (Cytivia).

Plasmids encoding the CR9114 Fab heavy and light chains for X-ray crystallography were dually transfected into Expi293 cells with the Fab heavy chain also encoding a Strep Tag II at the C terminus. Cell supernatant was harvested at day 5 when cells reached ∼80% viability, diafiltered to remove destinbiotin from the supernatant, then CR9114 Fab was purified using a StrepTrap HP column (GE Healthcare). The Strep Tag II was proteolytically cleaved using TEV protease (AcTEV protease, Thermo Fisher Scientific) prior to size exclusion chromatography in buffer containing 10 mM Tris pH 7.5, 150 mM NaCl.

#### Recombinant Hemagglutinin

Plasmids encoding for H1_NC99, H1_stem_NC99, H3_HK68, H5_VN05, V7_Sh07 were kindly provided by the NIH. All HAs contained a C-terminal T4 trimerization site, Avi-Tag and 6x His Tag. Modified versions as stem-epitope KO mutants, GCN4 trimerization sites and stem constructs were ordered as linear dsDNA inserts from Twist Bioscience and cloned into the VRC vector from NIH. All recombinant HAs carry the Y98F mutation in the receptor-binding domain. HAs were produced using the Expi293™ expression system from Thermo Fisher Scientific. Supernatant was collected 6 days post transfection, filtered and purified via Ni-NTA affinity or StrepTrap™ HP, for GCN4 HA versions, followed by size exclusion on a HiLoad® 16/600 Superdex® 200pg column on an ÄKTA™ pure system (cytivia).

#### Design nanoparticles

The sequences of the FI6-focused design_04 and stem-epitope design were cloned into a pHLsec vector with an N-terminal 6x His Tag and a C-terminal ferritin from *Helicobacter pylori* (GenBank ID: QAB33511.1). Designs and ferritin were connected by a GS linker containing one glycosylation site (GGSGGSGGSGGSNGTGGSGGS). Ferritin-design nanoparticles were produced using the Expi293™ expression system from Thermo Fisher Scientific. Supernatant was collected 6 days post transfection, filtered and purified via Ni-NTA affinity followed by size exclusion on a HiLoad® 16/600 Superose 6 pg column on an ÄKTA™ pure system (cytivia).

#### Surface plasmon resonance to measure binding affinities

SPR measurements were performed on a Biacore 8K (cytivia) with HBS-EP+ as running buffer (10 mM HEPES pH 7.4, 150 mM NaCl, 3 mM EDTA, 0.005% v/v Surfactant P20, GE Healthcare). Ligands were immobilized on a CM5 chip (cytivia #29104988) via amine coupling. Approximately 1000 response units (RU) of IgG were immobilized, and designed proteins were injected as analyte in two-fold serial dilutions. The flow rate was 30 µl/min with a contact time of 120 s followed by 800 s dissociation time. After each injection, the surface was regenerated using 0.1 M glycine at pH 2.5. Data were fitted using 1:1 Langmuir binding model within the Biacore 8K analysis software (cytivia #29310604).

#### Size exclusion chromatography multi-angle light scattering (SEC-MALS)

Size exclusion chromatography with an online multi-angle light scattering device (miniDAWN TREOS, Wyatt) was used to determine the oligomeric state and molecular weight for the protein in solution. Purified proteins were concentrated to 1 mg/ml in PBS (pH 7.4), and injected into a Superdex 75 300/10 GL column (cytivia) with a flow rate of 0.5 ml/min, and UV_280_ and light scattering signals were recorded. Molecular weight was determined using the ASTRA software (version 6.1, Wyatt).

#### Circular Dichroism

Far-UV circular dichroism spectra were measured using a Chirascan™ spectrometer (AppliedPhotophysics) in a 1-mm path-length cuvette. The protein samples were prepared in a 10 mM sodium phosphate buffer at a protein concentration between 20 and 50 µM. Wavelengths between 200 nm and 250 nm were recorded with a scanning speed of 20 nm min^−1^ and a response time of 0.125 secs. All spectra were averaged two times and corrected for buffer absorption. Temperature ramping melts were performed from 20 to 90°C with an increment of 2 °C/min. Thermal denaturation curves were plotted by the change of ellipticity at the global curve minimum to calculate the melting temperature (T_m_).

#### Next-generation sequencing

After sorting, yeast cells were grown in SDCAA medium, pelleted and plasmid DNA was extracted using Zymoprep Yeast Plasmid Miniprep II (Zymo Research) following the manufacturer’s instructions. The coding sequence of the designed variants was amplified using vector-specific primer pairs, Illumina sequencing adapters were attached using an additional overhang PCR, and PCR products were desalted on PCR purification columns (Qiaquick PCR purification kit, Qiagen). Next generation sequencing was performed using an Illumina MiSeq 2 × 150 bp paired end sequencing (300 cycles), yielding between 0.45-0.58 million reads/sample.

For bioinformatic analysis, sequences were translated in the correct reading frame, and enrichment values were computed for each sequence.

#### Bead assay

Anti-mouse IgG kappa beads (BD Biosciences) were coated with purified mouse anti-human IgG (BD Biosciences) to allow the beads to bind to any human IgG antibody. Anti-human IgG coated beads were washed with PBS + 0.1% BSA, then incubated with 1μg of different human monoclonal stem-specific antibodies for 20 min at 4°C followed by another wash step. Lastly, stem monoclonal antibody coated beads were incubated with 30 ng of biotinylated fi6-focused design 03 or stem-epitope design probes previously labelled with Streptavidin fluorophores for 20 min at 4°C followed by a last wash. Beads were measured in a LSRFortessa X-50 flow cytometer (BD Bioscience) and analyzed using FlowJo (TreeStar).

#### Labeling of sorting probes

HA proteins were expressed, purified, biotinylated and conjugated with different fluorophores as described previously (Joyce, M. G. et al. 2016, Whittle, J. R. R. et al. 2014, Andrews, S. F. et al. 2019). Briefly, purified HAs and immunogen designs were biotinylated on the AVI tag residue using BirA enzyme (Avidity) according to manufacturer’s protocol. Biotinylated HAs were labelled with Streptavidin conjugated with different fluorophores.

#### Human B cell sorting and sequencing

PBMCs from VRC310 clinical trial (Ledgerwood, J. E et al. 2011, Ledgerwood, J. E et al. 2013) (NCT# NCT01086657) were thawed, treated with Benzonase nuclease (Millipore Corp.) and washed with PBS. Cells were stained in PBS + 0.1% BSA with Aqua Fluorescent Reactive (Invitrogen), anti-IgM PercPCy5.5 (G20-127, BD Biosciences), anti-CD19 ECD (J3-119, Beckman Coulter), anti-CD38 AF700 (HIT2, eBioscience), anti-IgG BV421 (G18-145, BD Biosciences), anti-CD3 BV510 (OKT3, Biolegend), anti-CD14 BV510 (M5E2, Biolegend), anti-CD56 BV510 (HCD56, Biolegend), anti-CD27 BV605 (O327, Biolegend), anti-CD20 BV750 (2H7, BD Biosciences), anti-IgM BB700 (G20-127, BD Biosciences), anti-CD20 APCH7 (2H7, BD Biosciences), anti-IgG BUV395 (G18-145, BD Biosciences), anti-CD38 BUV661 (HIT2, BD Biosciences), anti-CD19 BUV805 (SH25C1, BD Biosciences), anti-IgD BV570 (1A6-2, BD Biosciences) and the different HA and immunogen design probes. Probes used were H5 HA A/Indonesia/05/2005, H3 HA A/Hongkong/2607/2019, fi6-focused design 03 and stem-epitope design. Cells were stained for 30 minutes at 4°C, washed with PBS + 0.1% BSA and analyzed on a FACS Aria II (BD Bioscicences) or FACS Symphony A5 (BD Biosciences) using Diva software. Cells were gated on live singlets CD3-CD14-CD56-CD19+ CD20+ CD38lo IgM-IgD-IgG+ memory B cells and all fi6-focused design 03 or stem-epitope design were single cell sorted into 96 well plates. Single sorted cells were submitted to a nested reverse transcriptase polymerase chain reaction (RT-PCR) to amplify the immunoglobulin heavy and light chains genes as described previously (Tiller et al. 2008). cDNA was made from dry-sorted plates using Superscript II Reverse Transcriptase (Thermo Fisher). Heavy, Kappa and Lambda chain genes were separately amplified by two rounds of PCR from the same cDNA using DreamTaq DNA polymerase (Thermo Fisher). All amplified products were sequenced by Genewiz and analyzed by IMGT. Heavy and light chain genes of interest were cloned into expression vectors by Genscript. 293Expi cells were transfected with the paired heavy and light chain plasmids using ExpiFectamine (Thermo Fisher), the antibodies were purified from the supernatant using a protein ASepharose (Cytiva) column and concentrated using a 30k amicon concenrator (Millipore).

#### Antigenicity assay

Monoclonal antibodies from B cells isolated using FI6-focused design_03 and stem-epitope design were tested against group 1 and 2 HAs through Meso Scale Discovery (MSD) technology. 386-well Streptavidin coated plates (MSD) were blocked with PBS + 5% MSD blocker A buffer (MSD). Plates were then washed with PBS + 0.05% Tween 20 (Thermo Fisher) and coated separately with 1 μg/ml of biotinylated FI6-focused design_03, stem-epitope design, H1 A/California/04/2009, H2 A/Canada/790/2005, H3 A/Wisconsin/67/2005, H5 HA A/Indonesia/05/2005, H7 HA A/Shanghai/02/2013, H9 HA A/Hongkong/1073/1999 and H10 HA A/Jiangxi-Donghu/346/2013. Plates were washed again, and monoclonal antibodies were added serially diluted, starting at 1μg/ml. Plates were washed and incubated with 1μg/ml of SULFO-Tag anti-human IgG detection antibody. After incubation, plates were washed and read with an MSD Imager 2400 machine using the MSD reading buffer. Data was analyzed using GrahPad Prism software.

#### Cell lines and viruses

MDCK-cells were cultured in DMEM (ThermoFisher Scientific) supplemented with 10% heat inactivated FBS (ThermoFisher Scientific) under 5% CO2 atmosphere at 37°C. MDCK-SIAT1 Were cultured as MDCK but with the addition of 500 µg/ml geneticin (Gibco). We employed the following viruses in this study: A/Puerto Rico/8/34 (H1N1), A/California/07/2009 (H1N1), A/HKx31 (H3N2), A/Michigan/45/2015(H1N1), A/Singapore/INFIMH-16-0019/2016 (H3N2) and A/Environment/Suzhou/SZ19/2014 (H7N9). Viruses were propagated in 10 days old embryonated chicken eggs (VALO BioMedia) or MDCK SIAT1 cells.

#### Serum binding to virus infected cells

The assay was performed as previously described (Angeletti et al., 2019). Briefly, MDCK cells were infected using A/Puerto Rico/8/34 (H1N1), A/California/07/2009 (H1N1), A/HKx31 (H3N2) viruses, at MOI = 3 for 5h at 37°C in DMEM medium with gentamycin (Thermo Fisher, Waltham, MA) and TPCK-Trypsin (BioNordika, Solna, Sweden). After incubation, cells were incubated with a serial dilution of sera for 60 min at room temperature. After washes with DPBS + 5% FBS + 0.5mM EDTA (Thermo Fisher, Waltham, MA), cells were stained with BV421 anti-mouse kappa light chain (clone 187.1) (BD biosciences, San Jose, CA) for 30 min at 4°C.

Cells were washed again and permeabilized using a fixation/permeabilization buffer from the Foxp3 / Transcription Factor Staining Buffer Set (Thermo Fisher, Waltham, MA) for either 30min or overnight at 4°C. Now, cells were stained with an AF488 coupled (AF488 Protein Labelling Kit, Thermo Fisher, Waltham, MA) anti-Influenza A virus NP (clone H16-L10-4R5 (HB65)) antibody (Bio X Cell, Lebanon, NH) in permeabilization buffer. The samples were acquired on a CytoFLEX flow cytometer (Beckman Coulter, Indianapolis, IN) and analyzed with the GraphPad Prism9 software.

#### ADCC assay

ADCC assay was performed as previously described (Kosik et al. 2019). Briefly, 10,000 MDCK cells were seeded into 96-well white flat bottom plates (Thermo Fisher, Waltham, MA) and left to settle overnight. The following day, cells were washed twice with DPBS (Thermo Fisher, Waltham, MA) and infected with following viruses at MOI=3 in RPMI 1640 medium with 4% ultralow-IgG serum (Thermo Fisher, Waltham, MA): A/Puerto Rico/8/34 (H1N1), A/California/07/2009 (H1N1) and A/HKx31 (H3N2). After 6 h, 25 µl of an 1:40 dilution of mouse sera were combined with 50,000 in 25 µl of effector cells (Jurkat cell line expressing luciferase gene under control of the NFAT response element and stably expressing FcγRIV; Promega) in RPMI 1640 medium with 4% ultralow-IgG serum (Thermo Fisher, Waltham, MA). The serum-effector cell mixture was added onto the MDCK cells (effector cells:target cells = 5:1). After overnight incubation at 37°C in 5% CO_2_, 50 µl of Bright-Glo Luciferase Assay lysis/substrate buffer (Promega) was added and luminescence was measured within 5 min using a SpectraMax® i3x plate reader (Molecular Devices, San Jose, CA). Fold induction was calculated over a no sera containing control using GraphPad Prism9 software.

#### Mouse immunizations

All animal experiments were approved by the Veterinary Authority of the Canton of Vaud (Switzerland) according to Swiss regulations of animal welfare (animal license VD3461). Female six-week-old BALB/c/cjRJ mice were purchased from Janvier Labs and acclimatized for one week. Hemagglutinins were used at a concentration of 20 µg/mL and design particles at 40 µg/mL. Immunogens were diluted with PBS (pH 7.4) to the intended concentration and mixed 1:1 with AS03 adjuvant right before the injection. Each mouse was injected intramuscularly in the hind leg with 50 µL, corresponding to 1 µg of HA and 2 µg of design particles. Immunizations were performed on day 0, 21 and 42. Homologous immunizations of stem-epitope particles were adjuvanted. Primed only immunizations with H1 (NC99) were adjuvanted, but H3 (HK68) primed only injections were not. Boost injections with stem-mimetic particles were also adjuvanted in the H1 primed scheme whereas H3 primed stem-mimetic boost injections were not. Tail bleedings (∼100 µL) were performed on day 0, 14 and 35. On day 56 mice were sacrificed, mice were anaesthetized with isoflurane and blood was drawn by cardiac puncture.

#### ELISA

Nunc MediSorp plates were coated with antigen (recombinant HA, design scaffolds or wildtype scaffolds) overnight at 4°C in PBS (pH 7.4). Plates were blocked with blocking buffer (PBS + 0.05% Tween + 5% skimmed milk powder (sigma, #70166) for 2h at room temperature (RT). Plates were washed 4 times with PBST (PBS + 0.05% Tween). Mouse sera was serially diluted in dilution buffer (PBS + 1% BSA) and incubated for 2h at RT. Plates were washed again 4 times with PBST. Anti-mouse-Fc HRP-conjugated antibody was diluted 1:5000 in dilution buffer and incubated 1h at RT. Plates were washed again 4 times and developed by adding 100 µL of TMB solution per well. The reaction was stopped after 5 minutes 100 µl with 0.5M HCl. The absorbance at 450 nm was measured on a Tecan Safire 2 plate reader, and the antigen specific titers were determined as the reciprocal of the serum dilution yielding a signal two-fold above the background.

#### Whole Virus ELISA

X31(H3N2) and California 0709 (H1N1) virus were UV-inactivated on ice for 30 min. 96-well plates (Greiner Bio-One GmbH, Kremsmünster, Austria) were then coated with either whole UV-inactivated virus or HAH1 purified from PR8 virus as previously described (Angeletti et al. 2019). After at least an overnight incubation at 4°C, plates were blocked with 5% BSA. Subsequently, after washing three times, sera to be tested were diluted in serial 2-fold dilutions down the plate. Plates were then incubated at 37°C for 1.5h. After another wash, goat anti mouse H+L chain IgG coupled to peroxidase (Vector Laboratories, Burlingame, CA) was used for staining. Plates were washed again and then developed with TMB (Thermo Fisher, Waltham, MA) and H_2_SO_4_. Plates were read at 405nm on a plate reader (Tecan, Männedorf, Switzerland).

#### Flow cytometry

Animals were immunized thrice with the stem design particle intra muscular into the left hind leg, three weeks apart between immunizations and one week after the last injection, mice were sacrificed. Iliac and Inguinal lymph nodes were pooled and analysed. Cells were stained in DPBS + 5% FBS + 0.5mM EDTA with anti-IgD BV786, clone 11-26c.2a (BD Biosciences, San Jose, CA), anti-CD3e BV510, clone 145-2C11 (BD Biosciences, San Jose, CA), anti-NK1.1 BV510, clone PK136 (Biolegend, San Diego, CA), anti-B220 APC-Cy7, clone RA3-6B2 (BD Biosciences, San Jose, CA), anti-GL7 PE-Cy7, clone GL7 (eBiosciences, San Diego, CA), anti-CD38 FITC, clone 90/CD38 (BD Biosciences, San Jose, CA), anti-CD19 BUV395, clone 1D3 (BD Biosciences, San Jose, CA), stem design mimetic conjugated to BV421 (BD Biosciences, San Jose, CA), recombinant HAH1 California 07/09 and recombinant HAH3 Brisbane (Protein Production Sweden, Gothenburg, Sweden) conjugated to APC and PE (Thermo Fisher, Waltham, MA) respectively. The labeled cells were run and data acquired using BD LSR Fortessa X-20 or BD FACSAria Fusion (BD Biosciences, San Jose, CA), data was analyzed using Flow-Jo software (BD Biosciences, San Jose, CA).

#### Crystallization and X-ray data collection

Immunogen designs were mixed with their corresponding Fabs at a molar ratio of 1:2, incubated overnight at 4°C and purified by sized exclusion chromatography in a low salt Tris buffer. Crystal setups for the FI6-focused_design-Fab complex were performed in MRC 2 Well UVXPO Crystallization Plate (Douglas Instruments) with 200 nl protein + 200 nl reservoir solution pipetted by an Oryx 8 (Douglas Instruments). FI6-focused_03 alone crystallized in 20% (w/V) PEG 4.000, 15% (V/V) 2-propanol and 100 mM sodium citrate pH 5.5 within 2 days. Crystals grew to 200 µm within three days. FI6-focused_03·FI6 complex crystallized in 13% (w/V) PEG 8.000, 20% (V/V) glycerol, 80 mM sodium cacodylate pH 6.5 and 160 mM calcium acetate after 1 day. Crystals grew to 500 µm within 2 days. FI6-focused_04·FI6 complex crystallized in 20% (w/V) PEG 3.350, 200 mM ammonium citrate pH 5.0 overnight. Crystals grew for 2 days to 200 µm. Crystal setups were 20% (V/V) ethylene glycol was added to these crystals mother liquor prior to flash freezing. All crystals were exposed to synchrotron radiation and diffraction datasets of 360° at 0.1° steps were recorded. The data was processed in XDS.

For the stem-mimetic-Fab complex sparse matrix crystals screens were set up at a 1:1 ratio of protein to buffer and crystals of the stem-mimetic_01-CR9114 Fab complex appeared within 21 days in buffer containing 0.2 M Ammonium citrate dibasic, 20% w/v PEG 3350. Crystals were cryo-protected in mother liquor supplemented with 20% ethylene glycol and shipped to the Advanced Photon Source (APS) at Argonne National Labs for X-ray data collection. X-rays diffracted to a maximum resolution of 2.7 Å and scaled in space-groups C2.

#### Structure determination, model building, and refinement of the FI6-focused scaffold

Structures were solved by Molecular Replacement (MR, phenix.phaser) (Adams et al. 2010). MR results were manually mutated to correct sequences in coot. Resulting structures were refined in phenix.refine with manual adjustments in Coot (Emsley et al. 2010). The final resolution cutoff was determined by paired refinement (Karplus et al. 2012). Residues or sidechains without clear electron density at 1 σ were removed. Structure refinement statistics are summarized in Supplementary Table S2.

#### Structure determination, model building, and refinement of the stem-epitope mimetic

Diffraction data were indexed and scaled with HKL2000 (Minor et al. 2006), and Phaser (McCoy et al. 2007) was used for molecular replacement using PDB 1YA9 as a search model for stem-mimetic_01, and PDB 4FQH as the search model for the CR9114 Fab. Iterative rounds of reciprocal space and real space refinement were carried out in Phenix (Adams et al. 2010) and Coot (Emsley et al. 2010). The final structure was refined to *R*_work_/*R*_free_ values of 17/22% (Supplementary Table S2).

