## supplementary_materials for "Computationally designed stem-epitope mimetics elicit broadly reactive antibodies"

[#]These authors contributed equally

### Supplementary Figures

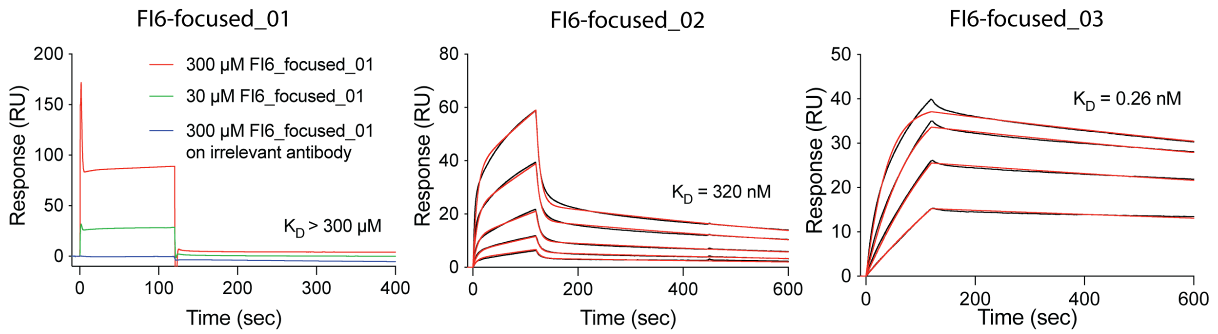


*Figure S1.* ***Affinities of FI6-focused intermediate designs to FI6 antibody measured by SPR.*** *Affinity of the initial FI6-focused_01 design was too low to determine a* $K_D$*. The irrelevant antibody used as negative control was an RSV specific antibody (D25). For the SPR measurements of FI6-focused_01 design FI6 IgG was immobilized on the chip and design was flown.For the SPR measurements of FI6-focused_02 and _03 design, designs were immobilized and FI6 Fab was added.*

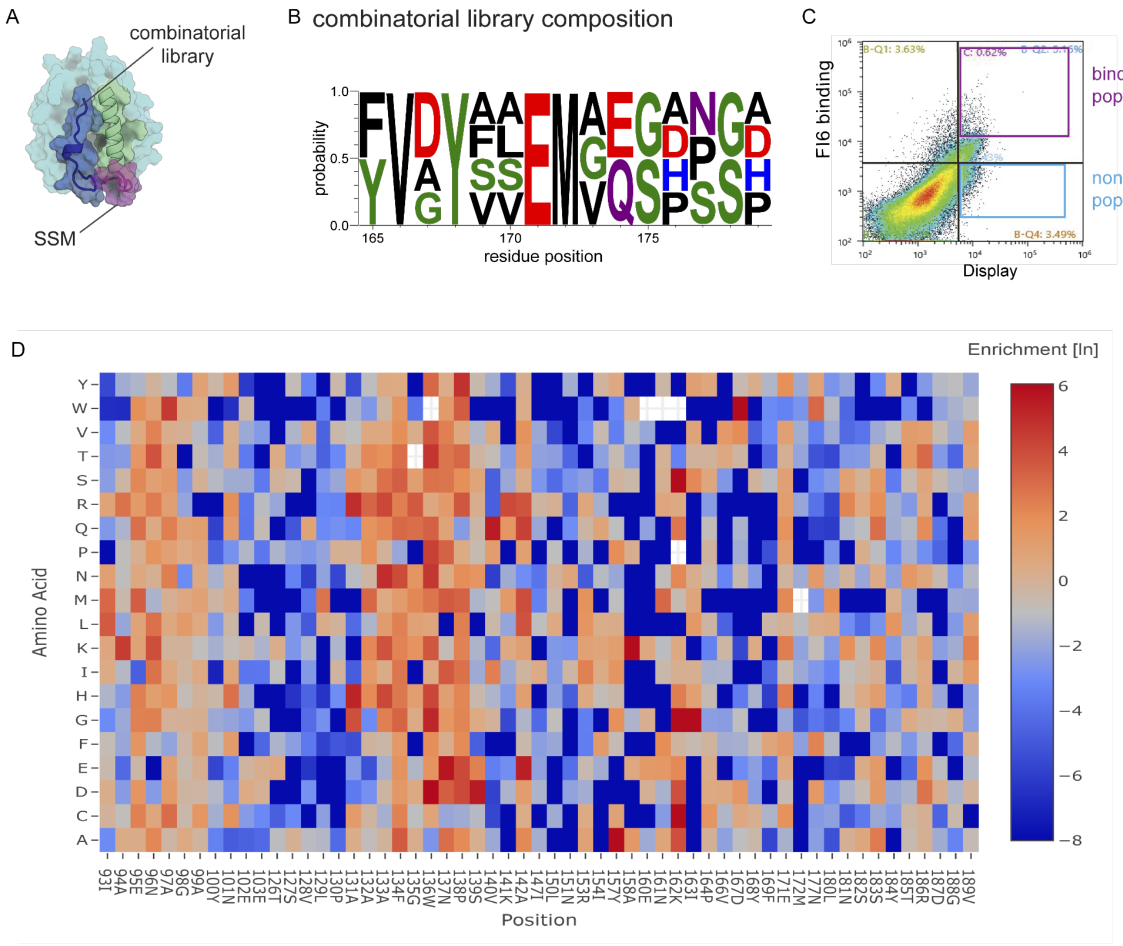


$$Enrichment = \ln\left(\frac{frequency\ in\ binding\ population}{frequency\ in\ non\ binding\ population}\right)$$

*Figure S2.* ***Overview of libraries for FI6-focused design affinity maturation. A.*** *FI6-focused design with targeted structural elements highlighted (green = grafted epitope helix, blue = hydrophobic pocket, purple = loop connecting the epitope helix to scaffold). The hydrophobic pocket was targeted with the combinatorial library. The connecting loop was targeted with the SSM library.* ***B.*** *Logo plot of positions targeted in combinatorial library. The library was sorted three times with decreasing concentrations of the FI6 antibody.* ***C.*** *Density plot of SSM library. Constructs binding strongly to the FI6 binding antibody and displaying on the yeast surface were sorted as binding population. Constructs without FI6 binding but displaying on yeast were sorted as non-binding population.* ***D.*** *Heat map of residues enriched in the binding population over the non-binding population.*

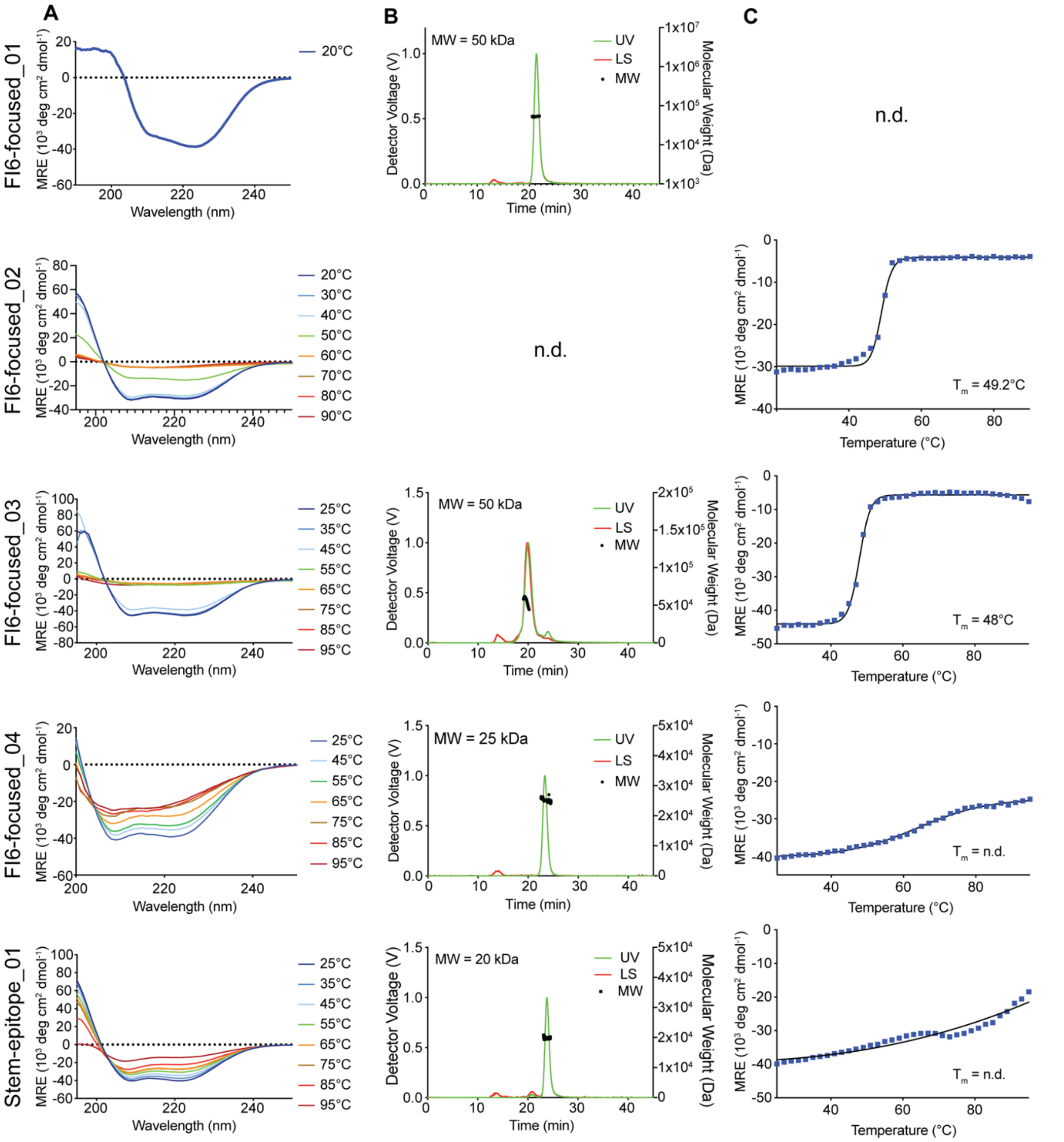


*Figure S3.* ***Biophysical characterization of relevant designs.*** *A. Circular dichroism (CD) spectra at indicated temperatures. B. Size exclusion chromatography coupled to multi-angle light scattering. C. Thermal denaturation curves. Designed proteins were melted from 20 to 90 °C by CD. The melting temperature ($T_m$) was determined by the change of ellipticity at the global curve minimum (208 nm).*

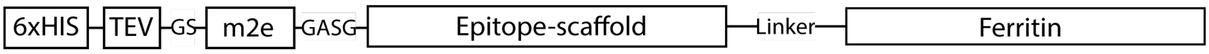


*Figure S4.* **Schematic overview of nanoparticle construct.** Epitope-scaffolds were fused C-terminally to ferritin separated by a GS-linker. They were labelled N-terminally with a His-tag for purification and a TEV cleavage site. The m2e T cell epitope was introduced between epitope-scaffold and His-tag.

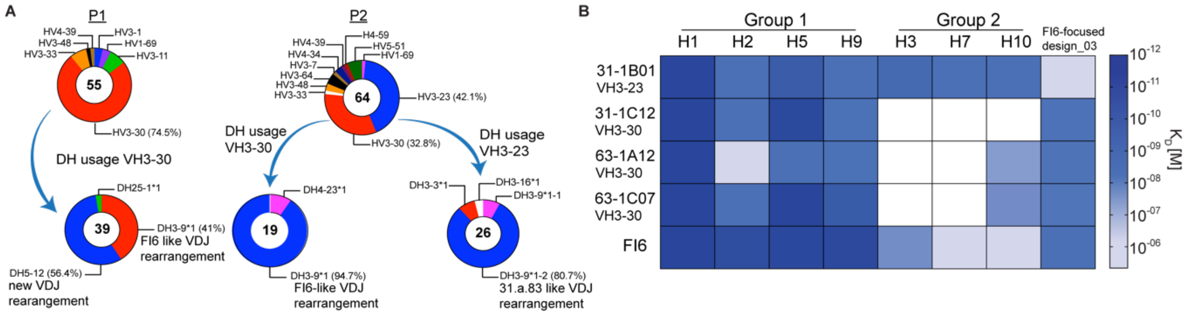


*Figure S5.* ***Analysis of human antibodies cross-reactive to FI6-focused design and H1. A.*** *Analysis of VDJ rearrangement pulled down antibodies and comparison to known bnAbs. B. Binding breadth and affinity of the different pulled down antibodies compared to FI6.*

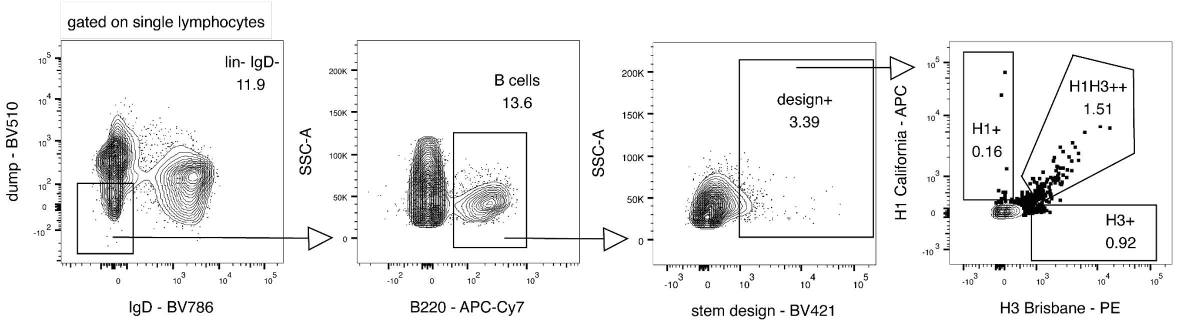


*Figure S6:* ***Gating strategy for B cell staining after immunization with 3x stem design. A.*** *Gating strategy to identify stem-design binding and H1-H3 cross-reactive B cells after immunization with 3x stem design. H1+H3+ cross-reactive B cells are selected as lin- IgD- B220+ design+.*

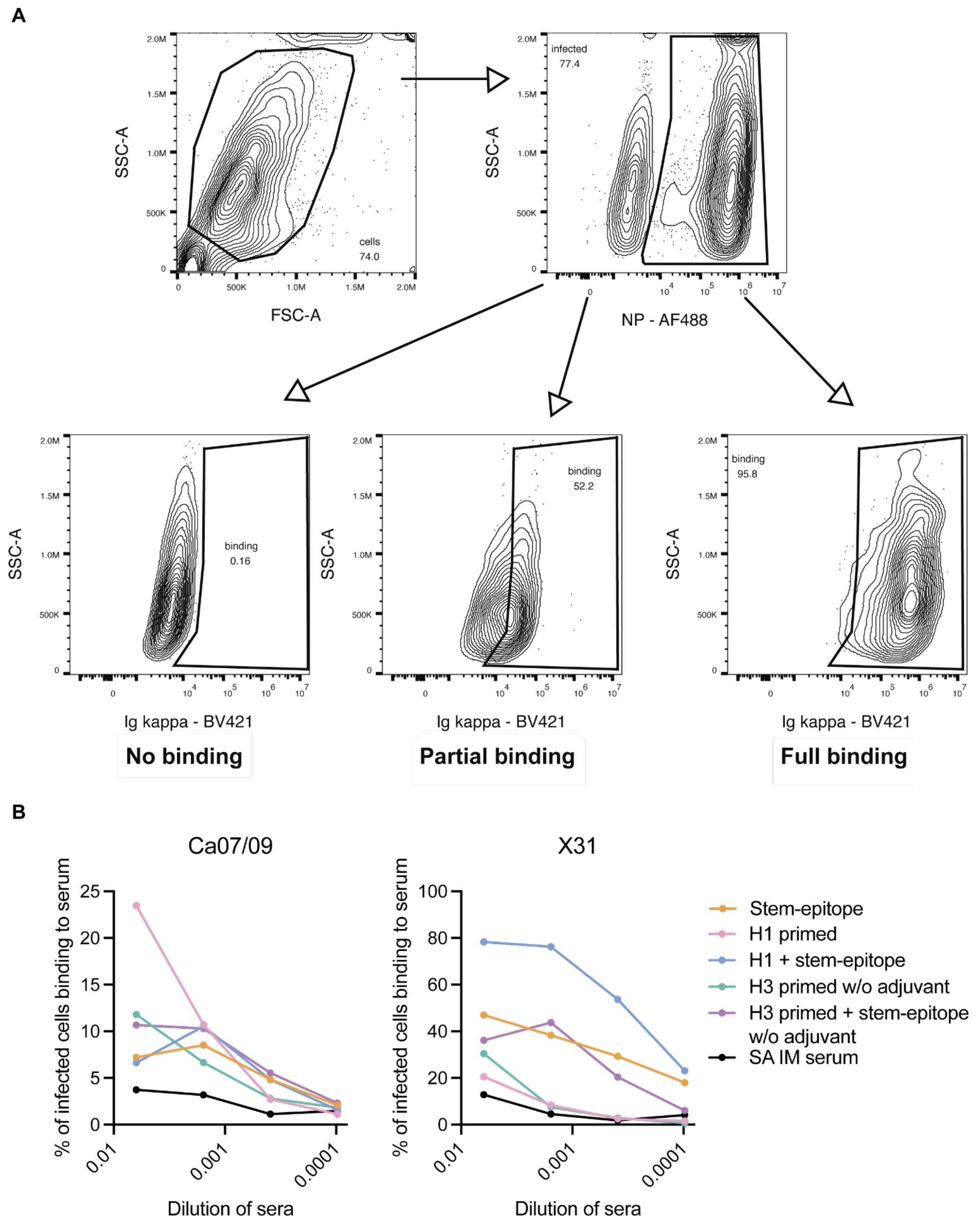


*Figure S7.* ***A.*** *Gating strategy to infected cells binding to serum. Gating strategy to identify serum binding to infected cells using flow cytometry. Analyzed are NP+ Ig kppa+ cells.* ***B.*** *Percentage of infected cells binding to serum ( antibody).* Frequency of infected cells binding to sera from immunized mice. Shown are 4-fold dilutions.

*Table S1.* ***Amino acid sequences of relevant designs and segments of the nanoparticle constructs.***

| Name | Sequence |
|---|---|
| m2e | SLLTEVETPIRNEWGSRSNDS |
| Linker | GGSGGSGGSGGSNGTGGSGGS |
| Ferritin | SKDIEKLLNEQVNKEMQSSNLYMSMSSWCYTHSLDGAGLFLFDHAAEEYEHAKK LIIFLNENNVPVQLTSISAPEHKFEGLTQIFQKAYEHEQHISESINNIVDHAIKSKDHA TFNFLQWYVAEQHEEEVLFKDILDKIELIGNENHGLYLADQYVKGIAKSRKS |
| FI6-focused design_01 | SDAQKQDWGNLKRYAEANKELVRKGKQKDRVVFMGNSITEGWVANDAAFFED NGYVGRGIGGQTSSHFLLRFREDVIKLAPALVVINAGTNDIAENAGAYNEEYTFGN IVSMVELARANKIKVILTSVLPAAAFGWNPSVKKATQAIIDLNTRIRNYAIENKIPYV DYFAEMVEGDNAALNSSYTRDGVHPTLEGYKVMEALIKKAIDKVL |
| FI6-focused design_02 | SDAQKQDWGNLKRYAEANKELVRKGKQKDRVVFMGNSITEGWVANDAAFFED NGYVGRGIGGQTSSHFLLRFREDVIKLAPALVVINAGTNDIAENAGAYNEEYTFGN IVSMVELARANKIKVILTSVLPAAAFGWNPSVKAATQAIINLNTRIINYAIENKIPFV DYFVEMAQSPNGDLNSSYTRDGVHPTLEGYKVMEALIKKAIDKVL |
| FI6-focused design_03 | SDAQKQDWGNLKRYAEANKELVRKGKQKDRVVFMGNSITEGWVANDAAFFED NGYVGRGIGGQTSSHFLLRFREDVIKLAPALVVINAGTNDIAENAGAYNEEYTFGN IVSMVELARANKIKVILTSVLPHAAAGWNESQKEATQAIINLNTRIINYAIENKIPFV DYFVEMAQSPNGDLNSSYTRDGVHPTLEGYKVMEALIKKAIDKVL |
| FI6-focused design_04 | SDAEKQDPANKKRYAEANKELVRKGKQKNRVVFMGNSITEGWVANDPAFFED NGYVGRGISGQTSSHMLERFEEDVIKLKPAVVVIMAGTNDIAENAGPYNEEQTFG NIVKMVELARAAKIKVILTSVLPHAAAPWNESQKEATQAIINLNTRIINYAIENKIPFV DYFVEMAQSPNGDLNSSYTRDGVHPNLEGYKVMEALIKKAIDKVL |
| Stem-epitope design_01 | GEPEVTDQLEWQSNQPWEQALNRFWDYLRWVQTLSDQVQEELQSSQVTQEL TALMEDTLTEAIAYMKELEEQLGPVAEETRLKLTQNVIDAITNLVNDMAELRNRL GQYRNEVHTMLGQSTEEIRARLSTHLRKMRKRLMRDAEDVQKALAVYKAGA |

*Table S2.* ***X-ray data collection and refinement statistics***

| | **Stem-epitope in complex with FI6 Fab** | **FI6-focused_03** | **FI6_focused_03 FI6** | **FI6-focused_04 FI6** |
|---|---|---|---|---|
| **PDB ID** | 8UZP | 9HI5 | 9HI4 | 9HI6 |

| | | | | |
|---|---|---|---|---|
| **Data Collection** | | | | |
| Wavelength (λ) | 1 | 0.9762 | 0.9801 | 1.037 |
| Resolution range (Å) | 37.79 – 2.71 (2.80 – 2.71) | 43.48 - 1.9 (1.968 - 1.9) | 44.84 - 1.7 (1.761 - 1.7) | 39.22 - 1.501 (1.555 - 1.501) |
| Space group | C2 | P 1 | P 1 21 1 | P 1 21 1 |
| Cell dimensions | | | | |
| *a, b, c* (Å) | 269.0, 55.9, 92.0 | 49.05, 63.31, 98.18 | 91.23, 78.14, 118.72 | 71.213, 47.869, 107.093 |
| α, β, γ (°) | 90, 90, 90 | 96.356, 99.528, 114.57 | 90, 100.601, 90 | 90, 106.126, 90 |
| Total reflections | 125034 (12471) | 280565 (28234) | 1250953 (127116) | 759197 (74564) |
| Unique reflections | 36153 (2434) | 79483 (7867) | 180128 (17937) | 110620 (10830) |
| Multiplicity | 3.5 (3.7) | 3.5 (3.6) | 6.9 (7.1) | 6.9 (6.9) |
| Completeness (%) | 90.7 (65.1) | 97.09 (95.43) | 99.88 (99.88) | 99.58 (98.19) |
| I/σI | 13.4 (2.9) | 12.38 (1.89) | 14.27 (1.08) | 11.78 (1.35) |
| Wilson B-factor | 44.60 | 30.57 | 27.18 | 19.50 |

| | | | | |
|---|---|---|---|---|
| R-merge | 0.142 (0.678) | 0.05935 (0.5936) | 0.07322 (1.642) | 0.09495 (1.155) |
| $CC_{1/2}$ | 0.81 (0.65) | 0.997 (0.752) | 0.999 (0.599) | 0.998 (0.595) |
| **Refinement** | | | | |
| Resolution (Å) | 37.79 – 2.71 | 43.48 - 1.9 | 44.84 - 1.7 | 39.22 - 1.501 |
| No. reflections | 34218 | 79457 (7865) | 180004 (17923) | 110607 (10829) |
| $R_{work}$ / $R_{free}$ (%) | 17/22 | 17/20 | 18/20 | 18/21 |
| No. atoms | | | | |
| Protein | 8684 | 6175 | 10033 | 4871 |
| Solvent | 27 | 639 | 1370 | 734 |
| R.m.s. deviations: | | | | |
| Bond lengths (Å) | 0.01 | 0.004 | 0.006 | 0.005 |
| Bond angles (°) | 1.64 | 0.64 | 0.82 | 0.78 |
| Ramachandran plot[#] | | | | |
| Favored (%) | 96.5 | 96.53 | 98.37 | 98.57 |

| | | | | |
|---|---|---|---|---|
| Allowed (%) | 3.5 | 3.47 | 1.63 | 1.43 |
| Outliers (%) | 0 | 0.00 | 0.00 | 0.00 |
| Rotamer outliers (%) | 6.34 | 0.34 | 0.28 | 0.00 |
| Clashscore | 11.30 | 1.66 | 1.38 | 2.53 |
| Average B-factor | 45.95 | 38.86 | 34.19 | 29.63 |
| Macromolecules | 45.98 | 38.53 | 32.97 | 28.65 |
| Solvent | 37.28 | 42.03 | 43.13 | 36.17 |

*R.m.s. deviation, root-mean square deviation.*
*Values in parentheses are for the highest resolution shell.*
*# Measured using Molprobity*
